# Deciphering eukaryotic *cis*-regulatory logic with 100 million random promoters

**DOI:** 10.1101/224907

**Authors:** Carl G. de Boer, Eeshit Dhaval Vaishnav, Ronen Sadeh, Esteban Luis Abeyta, Nir Friedman, Aviv Regev

## Abstract

Deciphering *cis*-regulation, the code by which transcription factors (TFs) interpret regulatory DNA sequence to control gene expression levels, is a long-standing challenge. Previous studies of native or engineered sequences have remained limited in scale. Here, we use random sequences as an alternative, allowing us to measure the expression output of over 100 million synthetic yeast promoters. Random sequences yield a broad range of reproducible expression levels, indicating that the fortuitous binding sites in random DNA are functional. From these data we learn models of transcriptional regulation that predict over 94% of the expression driven from independent test data and nearly 89% from sequences from yeast promoters. These models allow us to characterize the activity of TFs and their interactions with chromatin, and help refine *cis*-regulatory motifs. We find that strand, position, and helical face preferences of TFs are widespread and depend on interactions with neighboring chromatin. Such massive-throughput regulatory assays of random DNA provide the diverse examples necessary to learn complex models of *cis*-regulatory logic.

---

Understanding *cis*-regulatory logic would allow us to predict how gene expression is affected by changes to *cis*-regulatory sequences or regulatory proteins, providing both basic understanding of this fundamental process and helping determine the impact of genetic variants associated with common human traits and complex disease, most of which reside in regulatory sequences (reviewed in (*1*)). In general, learning models of *cis*-regulatory logic requires a training set of sequences and their associated expression levels. Analyzing natural regulatory sequences and the related gene expression profiles has met with some success (*2, 3*), but their limited diversity and shared homology mean that models are easily overfit (*3*), even when diversifying by mutagenesis (*4*). This is likely to be problematic in humans as well since genome-wide regulatory element assays identify fewer than 100,000 active elements in any given cell type (*5-7*). Measuring the expression of synthetic promoters, using either designed sequences (*8*) or designed elements (randomly-arranged (*9*)) can allow arbitrary hypothesis testing, but DNA synthesis technologies currently allow the creation of at most ~100,000 sequences. This limited scale means that TF binding sites (TFBSs) are often tested only in select affinities, contexts, positions, and orientations, leading to uncertain generalizability. Further, the sequences must be designed with specific hypotheses in mind and the hypotheses one can subsequently test are limited by this design. In contrast, the space of possible sequences or of combinatorial TF-TF interactions is vast. For example, testing all pairwise TF-TF interaction spacings only once each would require approximately 10^7^ sequences. Learning complex regulatory rules might require far more sequences than exist in the genome or have previously been assayed (*10*), such that predictive models of expression level from sequence alone remain elusive.

We hypothesized that random DNA offers a compelling alterative to learning regulatory models in eukaryotic genomes because the data, while not reflecting any specific organism’s regulatory DNA, could be generated at sufficient scale to learn complex models of gene regulation. Although TF motifs are expected to occur frequently by chance in random DNA (*11*), it is often tacitly assumed that *functional* TFBSs are rare: most TF motif instances are neither conserved nor bound, and it remains unclear whether TFBSs require additional factors to function (e.g. site clustering or interactions with neighboring factors) (*12*). Thus, it was not clear how often a random DNA sequence would include a sufficient number of TFBSs to drive reproducible expression levels, and whether a library of random sequence would span a sufficient dynamic range from which to uncover the rules of regulation. Past experiments that have used random DNA as a source of highly diverse sequences with which to study some aspects of gene regulation support our hypothesis. First, *in vitro* selection (or SELEX) relies on the fact that high-affinity TFBSs are present by chance in random DNA to select oligonucleotides from a random pool that are bound by a protein of interest (*13*) and use their sequences to define the specificities (*14*) and affinities (*15*) of TFs. Second, random DNA has also been used to diversify regions of native promoters (*16*), and to explore translational regulation (*17*) and to show that ~10% of random 100 bp sequences could serve as promoters in bacteria (*18*). However, no *in vivo* experiments have been conducted on the massive scale required to learn the complexities of *cis*-regulatory logic that can both (**1**) predict expression given any arbitrary sequence; and (**2**) explain how that sequence generated the expression level with interpretable features reflecting mechanisms of gene regulation.

To test our hypothesis, we developed the Gigantic Parallel Reporter Assay (GPRA) to measure the expression level associated with each of tens or hundreds of millions of *random* DNA sequences per experiment, and used these to learn models of *cis*-regulatory logic in the yeast, *Saccharomyces cerevisiae* (**Fig. 1** and **2**).

**Figure 1.**
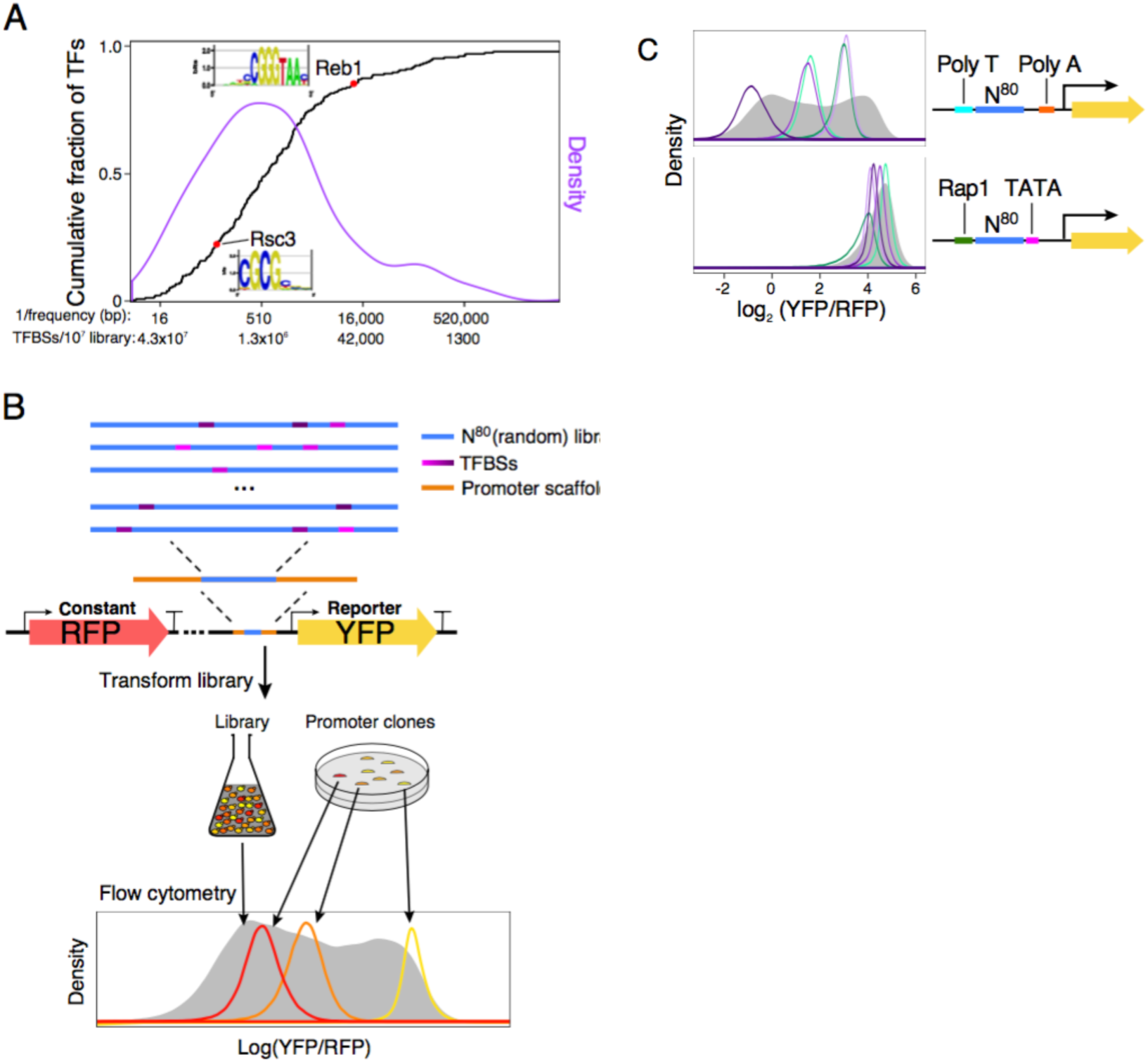
GPRA. (**A**) TFBSs are common in random DNA. Shown is the cumulative distribution function (CDF; black) and density (purple) of the expected frequency of yeast TF motifs in random DNA. The expected number of TFBSs in a library of 10^7^ random 80 bp promoters corresponding to each frequency is also indicated on the *x* axis. For instance, the relatively high information content (IC=14.59) yeast Reb1 motif is expected to occur on average once every 12,000 bp in random DNA, while Rsc3 (IC=7.78) should occur every 110 bp. (**B**) GPRA overview. From top: A library of random DNA sequences (N^80^ here, blue) is inserted within a promoter scaffold (orange) in front of a reporter (yellow arrow). By chance, the random sequences include many TFBSs (purple). When grown in yeast, the library would yield a broad distribution of expression levels (grey, bottom) as measured by flow cytometry, whereas each promoter clone would have a distinctive expression distribution (red, orange, yellow). (**C**) Random DNA yields diverse expression levels. For each promoter scaffold (right) shown are the expression distributions measured by flow cytometry (left) for the entire library (gray filled curves) and for a few selected clones, each from a different single promoter from each library (colored line curves).

**Figure 2.**
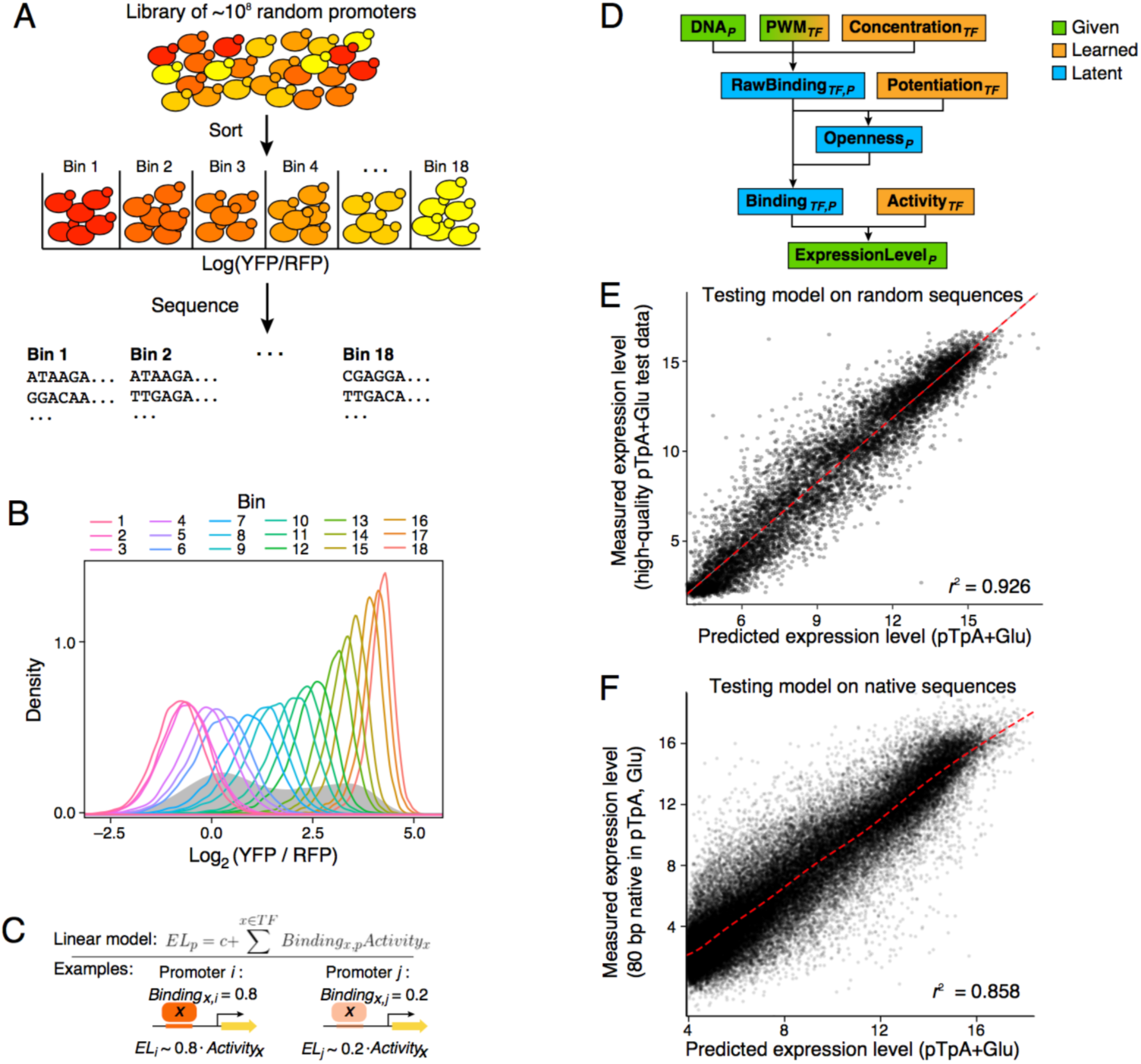
Expression models learned from a GPRA of 10^8^ random promoters are highly predictive. (**A**) Experimental strategy. Yeast GPRA library is sorted into 18 bins by the YFP/RFP ratio of the reporter (top) and the GPRA promoters in each bin are sequenced. (**B**) Reproducibility of expression levels. Expression distributions (log_2_(YFP/RFP)) for cells from each bin after sorting as in (A) (color code, top) which were regrown and reassayed by flow cytometry. Expression distribution maintains the initial bin ranking. (**C**) Linear model of TF activation. An increase in binding of a TF (*Binding*_*x,p*_) results in a proportional change in expression (*EL*_*p*_), scaled by the activity of that factor (*Activity*_*x*_). (**D**) Computational “billboard” model. Observed promoter DNA sequence (DNA_*p*_) are related to expression (ExpressionLevel_*p*_) by TF binding and activity. TF motifs (PWM_*TF*_) are provided as input, but can be refined and are used to calculate *K*_*d*_s for each potential TFBSs. Three parameters are learned per TF (orange): Concentration*TF* (the amount of binding to each TFBS given its *K*_*d*_), Potentiation*TF* (each TF’s ability to open/close chromatin), and Activity*TF* (each TF’s impact on transcription). Latent variables (blue) are calculated from the inputs and learned parameters. (**E,F**) Accurate prediction of expression from new random DNA and native yeast promoter sequences. Model-predicted expression (pTpA+Glu; *x* axis) *vs.* actual expression level (*y* axis) for (E) high-quality random 80 bp test data in the pTpA promoter scaffold grown in glucose and (F) native yeast promoter sequences, divided into 80 bp fragments and tested in the pTpA promoter scaffold grown in glucose.

First, we estimated that random sequence contains abundant yeast TFBSs. Consistent with previous models (*11*), TF motifs are expected to occur frequently in DNA uniformly sampled from the four bases (heretoafter “random DNA”) (**Fig. 1a, Methods**). For example, 58% of motifs are expected to occur at least once every 1,000 bp and 92% to occur at least once every 100,000 bp. Consequently, a random 80 bp section of DNA is expected to have on average ~138 yeast TFBS instances, comprised of partly overlapping sites for ~68 distinct factors. Thus, in a library of 10^7^ 80 bp random promoter sequences (as we create below), we expect, on average, that >90% of yeast TFs will have over 10,000 distinct instances of their respective TFBSs included, and most will have far more instances (**Fig. 1a**).

Next, we experimentally demonstrated that such random DNA yields diverse expression levels when used in a promoter library in yeast. To robustly quantify promoter activity (**Methods, Fig. 1b**) we used a previously described (*8*) episomal dual reporter system expressing a constitutive red fluorescent protein (RFP) and a variable yellow fluorescent protein (YFP), where log(YFP/RFP) measured using flow cytometry (**Methods**) reports a normalized expression signal that is integrated over several generations and controls for extrinsic noise (e.g. plasmid copy number and cell size) (*4, 19, 20*). We created ten synthetic promoter scaffolds and one based on a native promoter sequence (the *ANP1* promoter), each consisting of 50-80 bp of constant “scaffold” sequence on either side of a cloning site into which we inserted 80 bp of random DNA (–170 to –90, relative to the presumed TSS; **Fig. 1c and Supplemental Fig. 1, Methods**). In all instances, the random 80-mer libraries yielded diverse expression levels, up to a ~50 fold expression range, while individual promoter clones yielded distinct expression levels (**Fig. 1c** and **Supplemental Fig. 1, left**). When we tested a library where the scaffold sequences themselves were randomized (**Methods**), ~83% of random promoter sequences yielded measurable expression (**Supplemental Fig. 2**). This indicates that random DNA frequently contains functional TFBSs that modulate gene expression.

We then implemented GPRA as a robust assay, able to quantify the promoter activity of tens of millions of sequences in a single experiment. We created diverse libraries of ~10^8^ random promoters, transformed them into yeast, and sorted the cells by log(YFP/RFP) into 18 bins of equal intervals (**Fig. 2a, Methods**). We regrew the yeast from each bin, and measured their expression distributions by flow cytometry, observing excellent reproducibility (**Fig. 2b, Methods**). We sequenced the promoter libraries from each bin and used the distribution of reads for each promoter to estimate expression levels (**Methods**). Because the complexity of each promoter library (>10^8^) was greater than the number of sorted cells (<10^8^), most promoters (~78%) appear in only one bin, often representing one observation (read) from one cell containing that promoter and yielding a discrete expression level. While this leads to ~24% error in our expression estimates of individual promoters, the ability to create a vast number of examples with this approach outweighs this challenge, and yields highly informative data from which to learn rules of expression regulation, as we show below.

Altogether, we measured the expression output of 102,371,025 promoters with GPRA spanning two primary promoter libraries (each complexity > 10^8^) containing a random 80-mer with either: (**1**) an upstream poly-T sequence and downstream poly-A sequence (pTpA; **Fig. 1c**); or (**2**) an upstream Abf1 site and a downstream TATA box (Abf1TATA; **Supplemental Fig. 1**). We assayed both libraries with yeast using glucose as the carbon source, and the pTpA library was also assayed using either galactose or glycerol as alternate carbon sources. We sequenced 15-31 million unique promoters per experiment (<30% of sorted cells; <21% of the theoretical number of promoters in the libraries). Each library was sequenced to a depth of 50-155 million reads and did not reach saturation (**Supplemental Fig. 3**).

TF specific effects are captured well by GPRA. Even though each specific promoter sequence is typically associated with a single observed read, aggregating signal across the library reveals that there is a relationship between the strength of a TFBS and the observed level of expression. To identify TFBSs within each sequence, we used position weight matrices (PWMs) (*21*) for each yeast TF to scan each promoter sequence and estimate its predicted occupancy (*22*). Some TFBSs had a strong effect on expression but explained only a small percent of expression overall (*e.g.*, Abf1, a high IC motif which is relatively rare in random DNA, **Supplemental Fig. 4** left, Pearson’s *r* = 0.10), whereas others, including many zinc cluster monomeric motifs, correlated very strongly with overall expression (*e.g.*, Rsc30 *r*=0.57; **Supplemental Fig. 4**, middle). The sum of the individual motif effects (348%) is much greater than what we can explain with a simple linear model (~47%) across the TFBSs (**Fig. 2c**), suggesting that there is significant redundancy between motifs. Moreover, cases where related motifs have distinct behaviors (*e.g.*, Rsc30 and Ume6; **Supplemental Fig. 4**) indicate that examining TFBSs in isolation may be misleading.

Thus, as a more faithful joint model of TF activity, we pursued a “billboard model” (*23*) (**Fig. 2d**) that relates TF occupancy to expression with a linear model, but also captures how competition with nucleosomes and chromatin remodeling can alter TF occupancy (**Fig. 2d; Methods**). Since nucleosomes can potentially prevent TF binding (*24*), the model infers promoter accessibility, which is used to scale the predicted occupancy of each TF (*e.g.*, allowing for the possibility that a good TFBS will remain unbound by the TF if inaccessible). However, because some TFs can displace nucleosomes, we also learn how each TF can modulate the binding of other TFs (potentiation). Potentiation can be learned from cases where TFBSs alter expression only in the presence of another binding site that “potentiates” the first, which we assume is primarily driven by chromatin opening. Once parameters that capture each TF’s transcriptional activity, concentration, and ability to potentiate other factors are learned, we also refine its sequence specificity (**Fig. 2d**;**Methods**).

When trained on our GPRA data, these models impressively explained up to 92.6% of expression in independent, high quality test data (**Fig. 2e**). We learned a separate model for each of the four high-complexity promoter datasets: pTpA in glucose, galactose, and glycerol, and Abf1TATA in glucose. We tested each model by its ability to predict expression values measured for high-quality test data generated by assaying a smaller independent pTpA library of ~100,000 promoters in glucose. The pTpA+glucose model had the best predictive performance (*r*^2^ = 0.926, **Fig. 2e**), but the galactose- and glycerol-trained pTpA models performed nearly as well (*r*^2^ = 0.904 and 0.843, respectively). This indicates that the primary contributors to gene expression in the context of random DNA are not regulated by carbon source. Overall, a remarkably high proportion of random promoter expression is explained by a billboard model.

Even more remarkably, our models trained on random DNA data from GPRA, predicted over 85% of the expression driven by sequences from native yeast promoters (**Fig. 2f**). To this end, we segmented each yeast promoter into 80 bp fragments from −480 to the TSS, and assayed these in the pTpA promoter scaffold, in yeast grown in glucose. When we applied the pTpA+glucose billboard model, which was trained on random DNA, to the native yeast sequences, it predicted the observed expression with high correlation (Pearson *r*^2^=0.858, **Fig. 2f**). This shows the power of models trained on random DNA and indicates that non-billboard regulatory mechanisms are either not predominant in yeast promoters, or are context dependent.

A key aspect of our model is that it is biologically interpretable, such that we can assess not only the accuracy of the model’s predictions of expression level, but also mechanistic features, such as TF function or chromatin organization that underlie these predictions. For example, the models trained on expression levels of random DNA also accurately predict finer features, such as chromatin accessibility in the libraries themselves and in the yeast genome. First, there was a good correspondence between the model’s predicted nucleosome occupancy and the occupancy we experimentally measured by MNase-seq in the libraries (**Methods**; Spearman ρ = 0.54-0.55; **Fig. 3a** and **Supplemental Fig. 5**), comparable to the agreement between experimental replicates (**Fig. 3a**). Moreover, the pattern of nucleosome accessibility predicted by applying the models trained on random DNA to the sequences in the yeast genome agrees well with previously measured *in vivo* nucleosome occupancy in yeast (**Fig. 3b**; **Methods**). The models accurately predict the nucleosome free region and −1 and +1 nucleosomes, showing that the sequences and expression measurements generated by GPRA are of sufficient quality to correctly infer how TFs regulate chromatin structure, without ever directly measuring chromatin.

**Figure 3.**
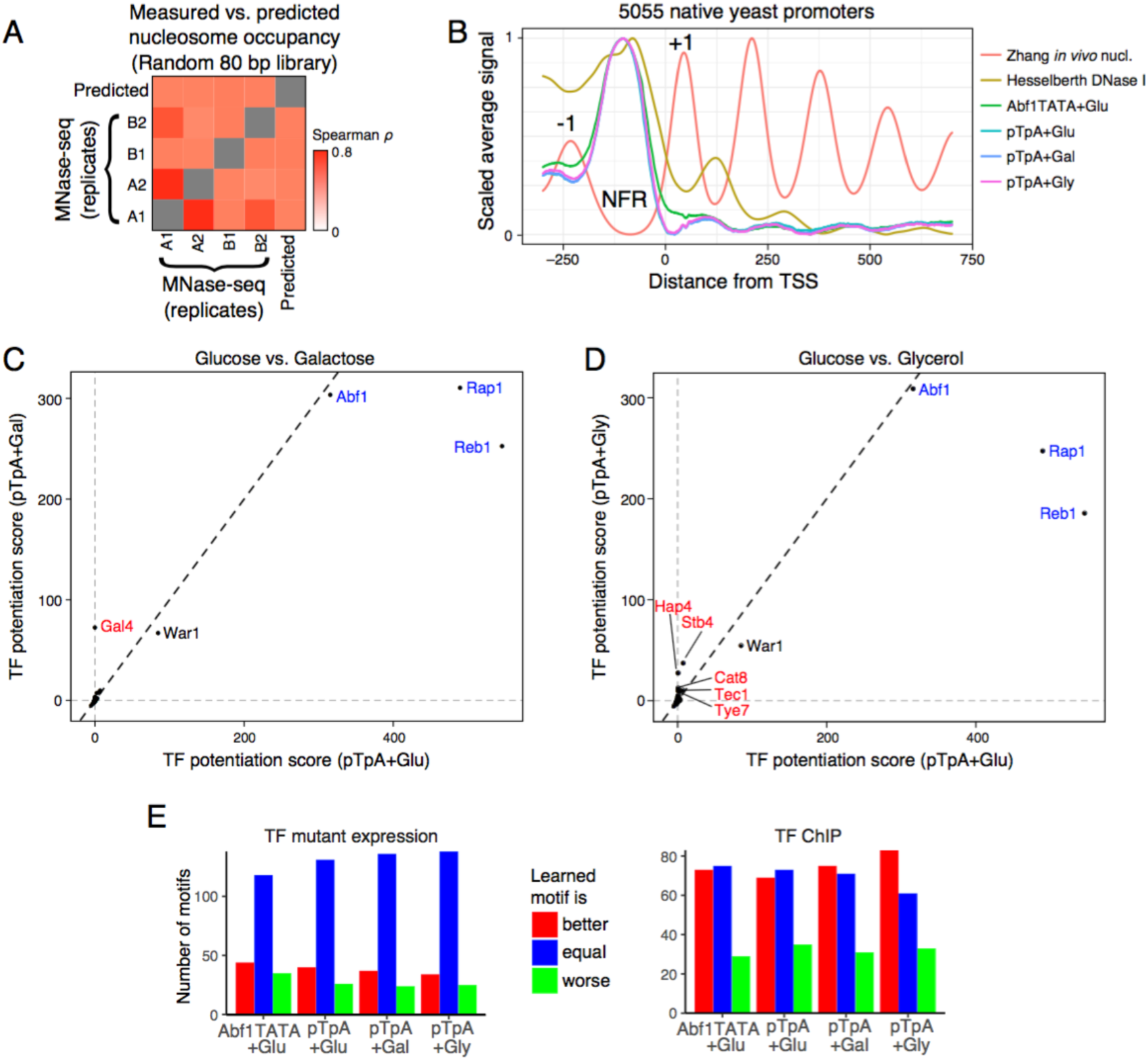
Billboard models learn biochemical activities of TFs. (**A,B**) Prediction of chromatin accessibility. (**A**) Pairwise Spearman correlations (color) between model-predicted nucleosome occupancy (1-predicted accessibility) and *in vivo* nucleosome occupancy measured by MNase (four replicates/conditions). (**B**) Metagene profile surrounding the TSS, based on *in vivo* nucleosome occupancy (Zhang (*25*)), DNase I hypersensitivity (representing accessibility; Hesselberth (*26*)), and model-predicted accessibility for each of the four billboard models. Each dataset is scaled. +1 and −1 nucleosome positions, and promoter Nucleosome Free Region (NFR) are indicated. (**C,D**) Prediction of chromatin opening ability. Shown is the predicted chromatin opening ability for each TF (dot) for pTpA models trained in glucose (*x* axes) *vs*. either (**C**) galactose or (**D**) glycerol (*y* axes). Blue: GRFs with known chromatin opening ability in all conditions; red: known and putative carbon source-specific regulators. (**E**) Motif refinement improves experimental predictions. The number of TFBS motifs (*y* axis) for which the model-refined motif predicted gene expression changes (TF mutant, top) or TF binding (ChIP, bottom) are better (red), worse (green), or equal (blue) to the original motifs, for each of the four models (*x* axis).

The models also accurately captured biochemical activities of TFs, including function as activators *vs*. repressors and as chromatin remodelers. The General Regulatory Factors (GRFs; Abf1, Reb1, and Rap1), which have known nucleosome displacing activity (*27-29*), were predicted to open chromatin by all models (positive potentiation scores) in all conditions tested (**Fig. 3c,d**). In addition, only in galactose, the galactose-specific regulator Gal4 was correctly (*30, 31*) predicted to open chromatin (**Fig. 3c**). TFs predicted to open chromatin only in glycerol included Hap4, Stb4, Cat8, Tec1, and Tye7 (**Fig. 3d**). There is strong support for these predictions: Hap4 is a global regulator of non-fermentative media like glycerol (*32*); Cat8 activates gluconeogenesis (*33, 34*) and Tye7 regulates glycolysis (*35*), which are the two endpoints of glycerol metabolism (*36*); Tec1 regulates pseudohyphal growth (*37, 38*), which is constitutive in glycerol (*39*); and Stb4 is predicted to regulate genes annotated for “oxidoreductase activity” (*21*), consistent with a role in using non-fermentable carbon sources. Moreover, the model-predicted activity for each TF weakly agreed with known activator/repressor status for models trained on glucose data (**Supplemental Fig. 6a**; hypergeometric P-values: 0.02 and 0.04), while there was no association for either galactose (P=0.34) or glycerol (P=0.79). However, TFs annotated as activators were predicted to have positive potentiation scores (*e.g.*, open chromatin) and TFs annotated as repressors had negative potentiation scores (*e.g.*, close it) for all models (**Methods**; hypergeometric P-values: 10^-3^ to 2×10^-5^; **Supplemental Fig. 6b**), consistent with open chromatin being more active. Thus, random sequences contain TFBSs in sufficient quantity to identify how and under what conditions they affect gene expression and chromatin, even for comparatively rare motifs (*e.g.*, those recognized by the GRFs).

Random DNA includes many TFBSs spanning a wide range of affinities for most TF. Consequently, the model was able to optimize the position weight matrices (PWMs) describing TF specificities, yielding refined motifs that have both better predictions and better performance on independent tasks. In particular, motif refinement (including by introducing additional bases of specificity) improved the predictive accuracy (*r*^2^) of the models on the independent high-quality test data by 9-12 percentage points. The four models often made the same changes to the motif, suggesting that the revised motif may more faithfully represent the true specificity of the factor (**Supplemental Fig. 7a**). Many of the refined motifs performed better than the original ones at the independent tasks of predicting which targets are bound by the cognate TF in the yeast genome by ChIP (*40*) and which yeast genes would change in expression when the cognate TF is perturbed (*41*) (**Fig. 3e**, **Supplemental Fig. 7b,c**, **Methods**). While many motifs were indistinguishable from the originals, of those that differed, the model-refinement improved the majority of motifs (**Fig. 3e**). Even though many of the original motifs were derived from this ChIP data (*21*), our refined motifs predicted the experiments better than the originals (**Fig. 3e**). This suggests that the refined motifs often more closely represent their cognate TF specificities.

Because motif position and orientation can affect TF function by modifying the TF’s ability to contact its biochemical target within mediator, the transcriptional pre-initiation complex (PIC), or surrounding chromatin, we next extended the billboard model with localized activity bias terms (**Methods**), allowing each TF to have a different activity for each possible binding position and orientation (**Fig. 4a**). This model must fit ~224 parameters per TF (instead of only four in the original model), including ~110 location-specific activity parameters for both strands of DNA (**Fig. 4b**), for a total of ~55,000 parameters in each model. Fitting such a model reliably with data of more traditional scale would not have been possible; however, the scale of our data means that promoter sequences still outnumber parameters ~360:1. Including these parameters significantly increased predictive value. For example, the extended pTpA+Glucose model now explained 94.3% (*vs*. 92.6%) of the expression measurements for the high-quality test data and 88.6% (*vs*. 85.8%) of the 80 bp native promoter sequences, representing ~20% decreases in the error in each case (**Supplemental Fig. 8**).

Our model shows that many TFs have strong position, orientation, and helical-face preferences (**Fig. 4b,c** and **Supplemental Fig. 9**). Predicted activators are often more active when binding distally within the promoter (*e.g.*, Abf1, Skn7, Mcm1; **Fig. 4b, Supplemental Fig. 9a,b**), while many predicted repressors were most repressive when binding proximal to the TSS (*e.g.* Ume6, Mot3; **Supplemental Fig. 9c,d**). Many TFBSs are strand-specific in their activity, often with a lower-than-expected activity distally, but for only one motif orientation (*e.g.*, Azf1, Mga1, Thi2; **Fig. 4b, Supplemental Fig. 9e,f**). Rarely, TFBSs are activating in one location and repressing in another (*e.g.,* Mga1; **Fig. 4b**). Some TFBSs showed strong periodicity along the length of the promoter (*e.g.*, Mcm1, Thi2, poly-A, Azf1; **Fig. 4b**, **Supplemental Fig. 9b,e,f**), consistent with a preference for a DNA helical face. Indeed, this appears to be widespread: comparing positional bias to a 10.5 bp sine wave, the correlations for each model were significantly higher than with randomized data (**Fig. 4c**; rank sum p<10^-120^; AUROC=0.84-0.87; **Supplemental Fig. 10**). Helical preferences tend to be strongest when TFBSs are proximal to the TSS (downstream of −150, relative the TSS). Since 150 bp is the approximate persistence length of dsDNA (*42*), this could reflect physical constraints of the promoter sequence: DNA is not flexible enough to allow efficient interactions between proximal TFs and the adjacent transcriptional pre-initiation complex, but after ~150 bp the DNA is flexible enough to relieve this effect. Notably, the positional biases sometimes varied for the same TFBS between models learned from different scaffolds (*e.g.*, Mga1, Skn7, poly-A, Mcm1; **Fig. 4b**, **Supplemental Fig. 9b**), suggesting that the positional preferences of TFs can depend on the surrounding context. Indeed, adding positional biases sometimes worsened the ability of models trained using one scaffold to predict expression for data generated in the other scaffold, but improved their performance on data generated using the same scaffold in another condition (**Supplemental Fig. 11**).

Overall, we showed that a massive-throughput, “big data” approach that measures the expression output of over 100 million random DNA sequences can provide a radically different scale compared to prior studies, surpassing the complexity of the human genome, and that this scale allows us to learn meaningful and complex models with remarkable predictive power. Using these data, we improved our ability to predict where TFs will bind, correctly predicted expression levels and chromatin organization, and identified factors that can remodel chromatin, including condition-specific regulators that can potentiate the activity of other TFs (**Fig. 4d**, top left). We showed that the majority of TFs have strand, location, and helical face preferences, demonstrating that *cis*-regulatory logic can be highly complex (**Fig. 4d**). We expect that more will be learned from this data: for example, a deep convolutional neural net trained on GPRA data explained ~98% of expression of our high-quality test data (E.D.V., unpublished results), representing a 60% reduction in the error. This exceptionally high-throughput approach allows researchers to determine the roles played the cell’s entire complement of TFs with a simple and inexpensive experiment.

**Figure 4.**
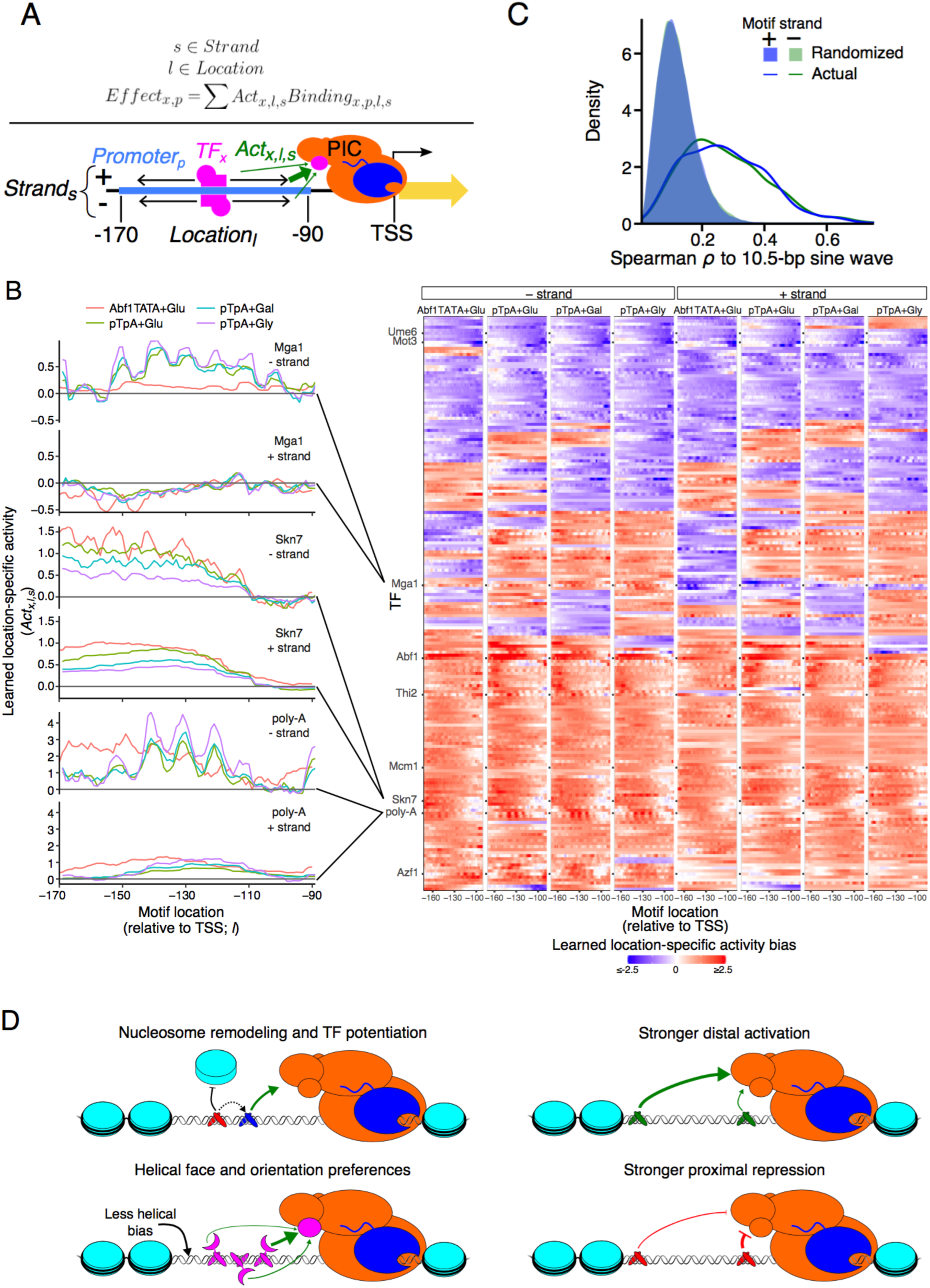
Position, orientation, and helical face preferences among yeast TFs. (**A**) Model with position and orientation-specific activities. For each TF, the model learns parameters for how much binding at each location within the promoter affects transcriptional activity. For example, this could reflect the TF’s ability contact the transcriptional pre-initiation complex (PIC). (**B**) Motif position and orientation effects on expression. Left: Each plot shows the learned activity parameter values (*y* axis) for motifs in each position (*x* axis) and strand orientation (upper and lower panels) for each model (colors). Right: Position-specific activity biases (color) for each TF (rows) at each position (columns) for minus (left half) and plus (right half) strand orientation for each of the four models (four subpanels). Only TFs for which all models retained the motif are shown. (**C**) Helical face preferences. Shown is the distribution of Spearman ρm between a 10.5 bp sine wave and the learned position-specific activity weights (as in **Supplemental Fig. 10**) for plus strand (blue line) and minus strand (green line) or with corresponding randomized data (blue and green shaded areas) for all four models. (**D**) Model of *cis*-regulatory logic. TFs display a variety of activity types. Some TFs potentiate the activity of other TFs by modulating nucleosome occupancy (upper left). Activators tend to have a greater effect on transcription when bound distally within the promoter (upper right), while repressors have the greatest effect when bound proximally (lower right). Many TFs show differential activity depending on the helical face or orientation of the TFBS, presumably through interaction with other factors bound nearby (lower left).

Our approach shows that using random DNA to study *cis*-regulatory logic *in vivo* is a highly accessible approach. We demonstrate that different random DNA sequences embedded within a scaffold can readily drive a broad range of expression levels and that random DNA has several major advantages for the study of *cis*-regulatory logic. The ease of generating massive libraries allows measurements of unprecedented scale, allowing us to learn complex models with very large numbers of parameters. While a library of designed sequences is limited to testing the hypotheses it was designed to test, a library of random DNA is useful for analyzing anything that occurs reasonably often by chance, even if they are not sufficiently common in the genome or where not planned for in advance. For instance, one could ask how G-quadruplex motifs affect expression (we saw no effect; data not shown). In addition, learning the effect of an element from thousands of examples with a wide range of affinities in thousands of unrelated contexts and in every relative position and orientation is likely to lead to more generalizable results than the “designed” approach of introducing a few element strengths into a few locations in a few common background sequences. Indeed, a “designed” approach will inadvertently introduce or destroy secondary TFBSs, which are hard to control for, and it can be difficult to partition the examples into training and test data due to due to shared design features or homology. Although using random DNA allows us to better learn how sequence dictates TF function, it necessitates joint modeling of the many variables that simultaneously affect expression.

The prevalence of functional TFBSs in random DNA and its demonstrated ability to modulate gene expression has implications for our understanding of the ways in which genes evolve. When a new gene is created by a mechanism like retrotransposition of an existing gene, the regulatory program, encoded by the DNA, must arise *de novo*. In bacteria, where there are no nucleosomes, random sequences have been shown to yield functioning promoters about 10% of the time (*18*). Here, we show that functional yeast promoter sequences also occur frequently by chance. In particular, ~83% of ~3,800 random promoter sequences (both random insert and random scaffold) appeared to be at least minimally active in glucose. Therefore, it may not be difficult to evolve basal gene regulatory sequences from previously non-regulatory DNA when a new gene is formed. Creating new enhancers in mammals may be similarly likely since mammalian TFs have, on average, even less specificity than those of yeast (*11*). This is also consistent with the observed fast evolutionary turnover of regulatory DNA, while overall expression programs are conserved (*43*), and with the hypothesis that many weak sites can impact transcription additively (*44*). Thus, newborn evolutionarily naive sequences will be primarily comprised of many weak TFBSs that have a comparatively weak effect on expression, potentially dominated by constitutive TFs with low specificity. Over evolutionary time, further mutations can optimize the specificity and effect of these new regulatory sequences.

In using GPRA, researchers will have to consider the scale needed for their question of interest. Since TFBS frequency depends on the specificity of the TF (**Fig. 1a**), more data will be needed to make inferences about rare TFBSs. In our analysis, the activity and potentiation parameters for each TF converged within the first 10% of the data. Conversely, an increase in data is important for refining or learning new motifs, and for finding position and orientation-specific activities. As noted above, since pairs of TFBSs are inherently rare in random DNA, learning all possible TF-TF interactions with GPRA, especially when considering competition (where both binding sites must be high-affinity), may require much bigger datasets. In principle, the GPRA approach could be applied to help decipher mammalian gene regulation. Although the increased complexity of having enhancers and more elaborate 3D genome topology may introduce challenges, such truly “big data” should allow learning models that explain how genetic variation affects gene expression and disease risk.

## Acknowledgments

We are grateful to Rani Nelken, Josh Weinstein, Atray Dixit, Brian Cleary, Karthik Shekhar, and Umut Eser for analysis advice; Christoph Muus, Brian Cleary, Atray Dixit, Yaara Oren, Thouis Jones, Luca Mariani, Karthik Shekhar, Justin B. Kinney, and David M. McCandlish for feedback on the manuscript; Toni Delorey, Jenna Pfiffner, and Caleb Bashor for experimental advice; Leslie Gaffney for help with figures; Patricia Rogers for cell sorting; and Eran Segal for the dual reporter yeast vector. CGD was supported by a Fellowship from the Canadian Institutes for Health Research and K99 award from the NIH. EDV was supported by the MIT Presidential Fellowship. Work was supported by the Klarman Cell Observatory at the Broad Institute, NHGRI Center of Excellence in Genome Science (CEGS), HHMI (AR), and the Israel Science Foundation ICORE on Chromatin and RNA in Gene Regulation (NF).

## Author Contributions

C.G.D. and A.R. drafted the manuscript, with all authors contributing. C.G.D. analyzed the data. C.G.D., E.D.V., E.A., and R.S. performed the experiments.

## Online Methods

### Theoretical TFBS abundance

We estimated the abundance of TFBSs in random DNA by analyzing the information contents (ICs) of known motifs associated with yeast TFs (*21*). The IC of a motif (*IC*_*motif*_) is proportional to the frequency (*f*_*motif*_) with which that motif is expected to be found on either strand of random DNA with the following relationship, where *IC*_*motif*_ is expressed in bits:

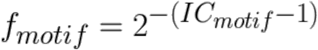

The number of instances present in a library of a given TFBS motif, assuming that binding sites are independent, is the number of positions in the library that could potentially contain a complete binding site multiplied by the expected frequency of the TFBS motif. For a library with a complexity of 10^7^, comprised of 80 bp sequences, the number of possible TFBSs is (80 – *length* _*motif*_ + 1) * 10^7^.

For **Fig. 1a**, we used the average motif length as the *length*_*motif*_ for all motifs so that the *x* axis could include frequency and the expected number of binding sites. For this analysis, motifs for zinc cluster monomers were excluded, since these are abundant in the database (*21*)and are likely to represent only a half TFBS. Several TFBS motifs that are long but generally have low IC content, were also excluded since they are unlikely to represent true TF specificities (**Supplementary Table 2**).

### Promoter library construction

For pTpA and Abf1TATA libraries, a single-stranded oligonucleotide pool was ordered from IDT containing the random 80 bp oligonucleotide flanked by arms complementary to the promoter scaffold for use with Gibson assembly. These oligonucleotides were double stranded with a complementary primer sequence and Phusion polymerase master mix (NEB), gel purified and cloned into the dual reporter vector, ensuring a complexity of at least 10^8^ for each library for libraries for which we measured expression, and 10^5^ for libraries for which we only inspected the overall expression distribution (**Fig. 1c** and **Supplemental Fig. 1**).

The two promoter scaffold sequences used for GPRA were:

For pTpA:

(poly-T; distal)

GCTAGCAGGAATGATGCAAAAGGTTCCCGATTCGAACTGCATTTTTTTCACATC

(poly-A; proximal)

GGTTACGGCTGTTTCTTAATTAAAAAAAGATAGAAAACATTAGGAGTGTAACACAAGACTTTCGGATCCTGAGCAGGCAAGATAAACGA (up to the theoretical TSS).

For Abf1TATA:

(Abf1 site; distal)

GCTAGCTGATTATGGTAACTCTATCGGACTTGAGGGATCACATTTCACGCAGTATAGTTC

(TATA-box; proximal)

GGTTTATTGTTTATAAAAATTAGTTTAAACTGTTGTATATTTTTTCATCTAACGGAACAATAGTAGGTTACGCTAGTTTGGATCCTGAGCAGGCAAGATAAACGA. In both cases, 80 Ns were inserted in between distal and proximal regions.

For the scaffold library (sequences in **Supplementary Table 1**), the library was cloned in two stages. In the first, the promoter scaffolds (synthesized by microarray synthesis) were amplified and cloned using Gibson Assembly. The resulting library had a common restriction site into which the N80 was cloned by ligation.

### Reporter assay

Libraries were transformed into yeast (strain Y8205 (*45*)) using the lithium acetate method (*46*), starting with 1L of yeast harvested at an OD of 0.3-0.4, ensuring at least 10^8^ cells were transformed (with the exception of the high-quality pTpA library, where a dilution series was performed to achieve the desired lower complexity). The yeast were then grown in SD-Ura for two days, diluting the media by 1:4 three times during this period. Media was then either changed to YPD, growing for at least 5 generations prior to cell sorting, or to YPGly and YPGal, with culture grown for at least 8 generations (due to the different carbon source). In the final 10 hours of growth prior to cell sorting, all cultures were allowed to grow continuously in log phase, never achieving an OD above 0.6, by diluting in fresh media. All cultures were grown in a shaker incubator, at 30°C and approximately 250 RPM.

Prior to sorting, yeast were spun down, washed once in ice-cold PBS, and then suspended in ice-cold PBS and kept on ice until cell sorting. Cells were sorted by log_2_(RFP/YFP) signal (using mCherry and GFP absorption/emission) on a Beckman-Coulter MoFlo Astrios, using the constitutive RFP under pTEF2 regulation to control for extrinsic noise. Cells were sorted into 18 uniform bins, done in three batches of six bins each, with the exception of the scaffold library, which was sorted into non-uniform bins to account for the higher variance at low expression levels and the larger dynamic range of the library. The FACS configuration varied between experiments (*e.g.*, different laser intensities), resulting in different baseline expression values. Post sort, cells were spun down and resuspended in SC-Ura (supplemented with 1% Gal for Gal sort), grown for 2-3 days, shaking at 30°C. The plasmids were then isolated, the promoter region amplified, Nextera adaptors and multiplexing indices added, and the resulting libraries sequenced with 76 bp, paired-end reads, using 150 cycle kits on an Illumina NextSeq sequencer, achieving complete coverage of the promoter, including overlap in the center. Libraries were not sequenced to saturation. For example, the pTpA+glucose experiment was sequenced with 155 million reads, yielding 31 million promoters, but doubling the number of reads is projected to only have yielded a further 8.5 million promoters (30%; **Supplemental Fig. 3**).

### Promoter sequence consolidation and expression level estimation

The paired end reads representing both sides of the promoter sequence were aligned using the overlapping sequence in the middle, constrained to have 40 (+/-15) bp of overlap for pTpA and Abf1TATA libraries and 16 (+/-10) bp for the scaffold library, and discarding any reads that failed to align well within these constraints. Note that only ~0.3μg of N80 DNA were received from IDT, and only ~10^8^ of these were successfully cloned; these are only a vanishingly small portion of the possible 4^80^ sequences in N80 (which would weigh ~10^26^ kg even with just one copy of each possible molecule). Thus, any very similar sequences we observe represent the same source promoter with high probability, with minor differences likely corresponding to PCR or sequencing errors. Consequently, promoters of pTpA and Abf1TATA libraries were aligned to themselves using Bowtie2 (version 2.2.1) (*47*) to identify clusters of related sequences, merging these clusters and taking the sequence with the most reads as the “true” promoter sequence for each cluster.

Expression levels for each promoter were estimated as the weighted average of bins in which the promoter was observed. For those observed only once, the expression level was the observed bin. The high-quality pTpA+glucose dataset, we restricted our analysis to only those ~10,000 promoters that had sufficient coverage (>100 reads each)

### Estimating the proportion of active random promoters

We also created a library of scaffolds that included 6204 scaffolds that were random but for the restriction site required to ligate in a random 80 mer, and the proximal 50 bp was ensured to be free of ATGs (to avoid out of frame reporter translation). Each scaffold included fixed distal and proximal promoter regions (-298:-195 and –103:-33, relative to the theoretical TSS, respectively) surrounding a variable 80 bp random oligonucleotide (-189:-109 regions). Each random scaffold was tested with ~660 random 80-mers, yielding approximately 4×10^6^ distinct random promoters total. This scaffold library was sequenced with a 300 cycle kit using a 190 bp read 1 and 112 bp read 2.

Promoters were first clustered into those sharing a common scaffold, using Bowtie2 to align to the known scaffold sequences (using the following parameters: -L 18 -p 4 -f --no-sq --no-head --np 0 --n-ceil C,100). Promoters were then sub-clustered within each scaffold using the sequences of the random 80-mers using CD-HIT (version 4.6.5, using the following parameters: -g 1 -p 1 -r 0 -c 0.96 -uS 0.05 -uL 0.05 -mismatch −1) (*48*), yielding a single consensus sequence for each promoter.

We estimated the proportion of random promoters that were expressed at detectable levels using the empirical log(YFP/RFP) distributions of regrown, previously-sorted, cells (as in **Fig. 2b**). We considered any bin above the lowest expression bin to be “expressed”, but since some cells might end up in this lowest expression bin upon re-sorting, we attempted to estimate the number of cells that would remain expressed upon resorting. AUROC statistics were calculated to estimate how well the cells sorted into each bin can be distinguished from those sorted into the not-expressed bin. Here, each AUROC is equivalent to the probability that a cell sorted into the corresponding expressing bin is expressed higher than a randomly selected cell from the not-expressed bin. Thus, cell proportions in expressing bins were weighted by the corresponding AUROC for that bin to get an estimate of the number of expressing random promoters, 83%.

### Testing native yeast promoters by GPRA

To test native yeast promoters in the GPRA system, the promoter sequences from the S288C reference genome (v64) were segmented into 80 bp fragments, overlapping by 40 bp, for a total of 11 fragments per promoter and 62,897 promoter fragments overall. We also included 8,027 promoters originally assayed within the high-quality pTpA N80 glucose experiment for use as controls (these were excluded from analyses evaluating model performance on native promoter sequences). The sequences were created by pooled oligonucleotide synthesis, including ends complementary to the pTpA scaffold. The fragments were amplified by PCR and cloned into the pTpA vector by Gibson assembly. The resulting library was transformed into yeast (S288C *ura3*Δ) and assayed as described above, with two replicates. We combined the two replicates, which showed some non-linearities resulting from differences in FACS binning procedures, using loess regression (span=0.1) to remove the non-linear relationship between one replicate and the average of the two replicates. After combining the replicates, the Pearson *r*^2^ between expression measurements in the combined replicates and the expression values originally measured for these promoters (from the high-quality pTpA N80 glucose experiment) was 0.977.

### Linear transcription model

TF motifs (**Supplementary Table 2**) were taken from the YeTFaSCo database (*21*) and supplemented with the poly-A motif (AAAAA), which we initialized to 100% A at all five positions. Motifs were trimmed to fill 30 bp 1-d convolutional filters, centering the motif if it was less than 30 bp, and, where motifs were longer than 30 bp, trimming off the least informative bases until it was 30 bp.

To identify dissociation constants, *K*_*d*_, for each TFBS motif and each potential binding site instance, motif filters were applied to DNA sequences of each promoter (*DNA*_*P*_) and their reverse complements by scanning them with the TFBS motif position weight matrix for each yeast TF (*PWM*_*TF*_). Binding to each site in the DNA was determined by the GOMER method using a fixed [*TF*_*x*_] that corresponds to the minimum *K*_*d*_ possible with the motif (and therefore a perfect match corresponds to 50% occupancy) (*22*). We considered all TFBSs, such that weak sites can also be influential, creating an affinity landscape for each TF across the region (*49*), and summed the predicted occupancy at each site, to obtain the expected occupancy for each TF of each sequence.

The expected binding (sum of all binding to all binding sites), assuming Michaelis-Menten equilibrium binding occupancies for all possible binding sites (location *l*, strand *s*) for TF *x* in promoter *p*, where *K*_*d*_s for each binding site are calculated from the position weight matrix:

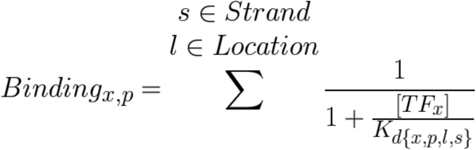

Correlations between predicted occupancy for each individual TF and expression level were done using these values (*Binding*_*x,p*_). We optimized a single weight for each TF (*Activity*_*x*_), representing the ability of that TF to activate or repress transcription.

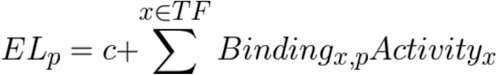

This model was implemented in Tensorflow, as described for the other models below, but without a regularization term.

### Billboard model of transcription

The billboard model includes parameters for TF concentration ([*TF*_*x*_]), TF activity (*Act*_*x*_), TF potentiation (*Pot*_*x*_), and TF activity limits (*AL*_*x*_). Motifs were trimmed, as before, but filling 25 bp 1-d convolutional filters. As described above, we use these filters, the DNA sequence of each promoter (*DNA*_*P*_), and the (now learned) TF concentration parameter to gain an initial estimate for TF binding that does not yet consider chromatin state, here called Raw Binding (*RB*_*x,p*_).

Some TFs can displace nucleosomes, so the model learns TF-specific parameters that capture the ability of each TF to modulate the binding of other TFs (*Pot*_*TF*_), which we assume is primarily driven by chromatin opening. Promoter accessibility is estimated as a logistic function on the potentiation-weighted *RB*_*TF,P*_ estimates, yielding a probability of the DNA being accessible (*Open*_*p*_):

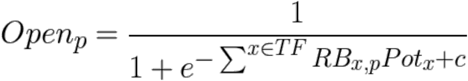

Since nucleosomes can potentially prevent TF binding (*24*), the previous estimate of binding (*RB*_*TF,P*_) is then scaled with this value, yielding the expected binding of each TF to each promoter (*Binding*_*TF,B*_):

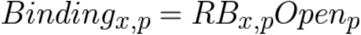

Because our promoters are small, we can reasonably assume that a TF that opens chromatin would open it for the entire 80-bp variable region: if the promoter is open, all TFs can bind unimpeded; if the promoter is closed, no TFs can bind. For example, a promoter that is predicted to be 0% accessible will have no TF binding, regardless of the TFBSs present in the sequence (*Binding*_*TF,P*_ = 0), while a promoter that is 100% accessible will have occupancy unchanged (*Binding*_*TF,P*_ = *RawBinding*_*TF,P*_). Thus, the model learns which TFs may, for example, open and close chromatin by their ability to potentiate the activity of other TFs (*i.e.*, TFBSs for TFs that affect transcription, but cannot open chromatin, only have an effect when “potentiated” by another factor, presumably by opening chromatin and allowing binding).

Finally, the predicted expression level (*EL*_*p*_) is the sum of binding values for each TF *x*, weighted by their learned effect on expression (*Act*_*x*_):

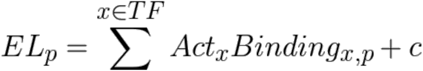

Here, the measured and predicted expression levels are in log space, corresponding to the log-space bins of YFP/RFP. One possible interpretation of the formulation above is that TF activities are proportional to how much the TF affects the zero-order rate constants for different steps of mRNA production, which would be multiplicative in linear space or additive (as above) in log space.

When activity limits for TFs (*AL*_*x*_) were included as a learned parameter, the expression level was instead calculated as follows, putting an upper limit on TF activity:

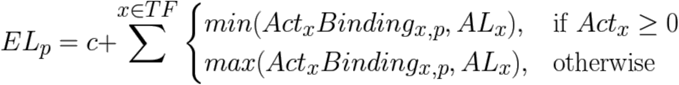

### Position-specific activity model

Position-specific activity models were built as an extension of the billboard model that included binding limits. Here, each potential TFBS position was allowed its own (learned) activity parameter. Position-specific occupancy was estimated similarly to before, but accounting for the strand (*s*) and binding location (*l*) of each TF (*x*) to each promoter (*p*):

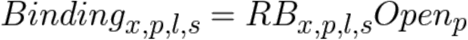

The transcriptional effect of each TF on each promoter (*Effect*_*x,p*_) was estimated using the position-specific activity parameters (*Act*_*x,l,s*_), which were implemented as a local scale of the overall TF activity (*Act*_*x*_):

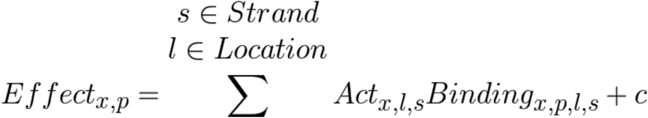

We then re-implement the binding limits as follows:

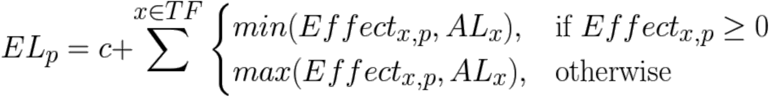

### Model learning

Parameters were learned iteratively, first learning TF activity and potentiation, then TF concentration, then allowing the motifs themselves to be changed, then including a parameter that limited the maximum binding of each TF, and finally learning position-specific activity parameters, each time, learning the new parameters and updating those previously included with a single pass through the data.

Transcriptional models were implemented in Tensorflow (*50*), minimizing the mean squared error between predicted and measured expression level using the AdamOptimizer and learning in batches of 1,024 promoters. In all cases (except the linear model above), potentiation and activity parameters were regularized with an *L*1 penalty (0.00001), motifs were regularized with an *L*2 penalty (0.000001), and position-specific activity biases (when present) were regularized with an L2 penalty (0.00001) on the difference between adjacent (by location *l*) activity biases. Learning rate was set to 0.04 for the epoch learning activity and potentiation parameters, 0.01 when also learning concentration, and 0.001 when also learning motifs, activity limits, and position-specific activities. All analyses used the models that did not include activity limits, with the exception of the comparisons to Miller et al. (*51*) data, and the position-specific activity model.

### Applying models to native sequences

Since the models above were designed to operate on relatively short sequences (~110 bp), scanning the yeast genome (R64) was done in tiling windows of 110 bp each, spaced at 1 bp intervals, yielding expression and accessibility predictions for nearly all bases in the genome.

To compare to chromatin organization in core promoters, the accessibility predictions were averaged across all yeast promoter sequences to yield a metagene plot, as was done for DNase (*26*) and nucleosome occupancy (*25*) data.

Predictions on the 80 bp fragments of native promoters tested in the pTpA scaffold were done as with other pTpA-scaffold model predictions.

### Comparing refined and original motifs

The original and model-refined motifs were evaluated for their ability to predict independent ChIP binding and TF mutant gene expression data. The GOMER method (*22*) was used to get a predicted binding occupancy of each sequence for the original and model-refined motifs. For ChIP data (*40*), ChIP-chip probes were scanned with the motifs, and their ability to predict ChIP binding for the corresponding TF was evaluated. For TF perturbation experiments (*21, 41*) promoter sequences were scanned with motifs, and their ability to predict expression changes when the cognate TF is perturbed (mutated, over-expressed, or deleted) was evaluated. In both cases, there were often multiple experiments for the same TF. We repeatedly sampled the data from each experiment (50% of the data sampled randomly 100 times, without replacement), and with each sample calculated the Pearson correlation coefficient between motif-predicted binding and biological measurement (gene expression, ChIP intensity) for both model-refined and original motifs. If the model-refined motif had a Pearson *r*^2^ greater than the original in at least 95% of samples, we considered the experiment to be predicted better by the refined motif. Conversely, if the original motif was better in at least 95% of samples, the experiment was considered to be predicted worse by the refined motif. A model-refined motif was considered to be better than the original if at least one experiment was predicted better and no experiment was predicted worse, while it was considered worse if at least one experiment was predicted worse and no experiment was predicted better. In all other cases, the motifs were considered equal. Motifs that were regularized out of the model (*i.e.* became neutral PWMs) were not considered in this analysis.

### Classifying TFs into activators and repressors by GO annotation

GO terms for yeast genes were downloaded from SGD (*52*) on Jan. 14, 2017. TFs annotated with a term containing any of “positive regulation of transcription”, “transcriptional activator”, “activating transcription factor binding”, or “positive regulation of RNA polymerase II” were labeled as activators. TFs annotated with “negative regulation of transcription”, “transcriptional repressor”, “repressing transcription factor binding”, or “negative regulation of RNA polymerase II” were labeled as repressors. Any annotated as both or neither were ignored for the purposes of testing for enrichment.

The models predicted that, most TFs opened rather than closed chromatin (*i.e.*, had positive potentiation scores; 64-66%) and most were predicted activators rather than repressors (53-55%), although most TFs in all four experiments were predicted to have little activity, consistent with many TFs being inactive in rich media (*53*).

### Promoter library MNase-Seq

Aliquots of the pTpA library, expected to correspond to ~100,000 (sample A) or ~200,000 (sample B) viable cells were each cultured in duplicate (Rep 1 and 2) in YPD for ~16 hours to an OD of ~0.4-1.0. For each sample, 0.5 mL of culture was pelleted and frozen to prepare input genomic DNA, and 3 mL of culture was crosslinked with 1% formaldehyde, washed twice with 1mL H_2_O supplemented with a protease inhibitor cocktail, and the pellet frozen for MNase treatment. These pellets were next spheroplasted using zymolyase, and spheroplasts were lysed in NP buffer (10 mM Tris pH 7.4, 50 mM NaCl, 5 mM MgCl_2_, 1 mM CaCl_2_, and 0.075% NP-40, freshly supplemented with 1 mM β-mercaptoethanol, 500 μM spermidine, and EDTA-free protease inhibitor cocktail) at a concentration of 2*10^6^ cells/ μl of NP buffer. 0.125 units of Worthington MNase were added per 10μl of lysed spheroplasts and MNase digestion was performed at 37°C for 20 minutes. MNase digestion was stopped by addition of equal volume of 2X MNase Stop Buffer (220 mM NaCl, 0.2% SDS, 0.2% sodium deoxycholate, 10 mM EDTA, 2% Triton X-100, EDTA-free protease inhibitor cocktail). MNased chromatin samples were treated with RNase A and proteinase K, reverse cross linked, separated on a 4% agarose gel and mononucleosome bands were isolated. Genomic DNA was prepared using the Masterpure Yeast Genomic DNA Preparation Kit (Epicenter). For both MNase and genomic DNA, the variable region of the promoter library was amplified, and adaptors added for sequencing using an Illumina NextSeq with 76 bp single-end reads.

Sequencing reads were mapped to all known promoters in any pTpA library using Bowtie2 (*47*). Only promoters with at least 20 reads in the input DNA and 1 read in the MNase data were kept for subsequent analysis. Input and MNase counts were scaled within each sample to yield counts per million (CPM) per promoter and the log ratio of MNase to input was compared between replicates and to the model’s predicted occupancy, corresponding to log(1-predicted accessibility). To combine MNase replicates, the log ratio of MNase to input was averaged for promoters present in both samples – those in only one sample were ignored. Similarly, pairwise correlations between samples in **Fig. 3a** reflect only the promoters common to both samples, and all promoters within the sample when comparing to the model’s predictions.

### Position and orientation-specific TF activities

In order to identify the approximate fraction of TFs displaying a 10.5 bp helical activity bias, the position-specific activities across the variable promoter region were compared to a 10.5 bp sine wave. First, we regressed out the overall positional activity bias using loess regression (span=0.5; green curves in **Supplemental Fig. 10**). These long-range trends were subtracted from the data, leaving only the short-range trends (blue curves in **Supplemental Fig. 10**), which were then compared to a 10.5 bp sine wave for 100 possible alignments of the sine wave, taking the largest magnitude correlation for each TF and strand, and calculating Spearman’s correlation coefficient, ρm. As background, the same procedure was performed after first shuffling the position-specific activity biases for 100 permutations of the data per TF. A P-value and AUROC were calculated describing the difference between the randomized and actual data for each model using Wilcoxon’s rank sum test.

### Data and software availability

Data are available at NCBI’s GEO: GSE104903, GSE104878. Open source code for our transcriptional models is available at https://github.com/Carldeboer/CisRegModels

## Supplemental Figures

**Supplemental Figure 1.**
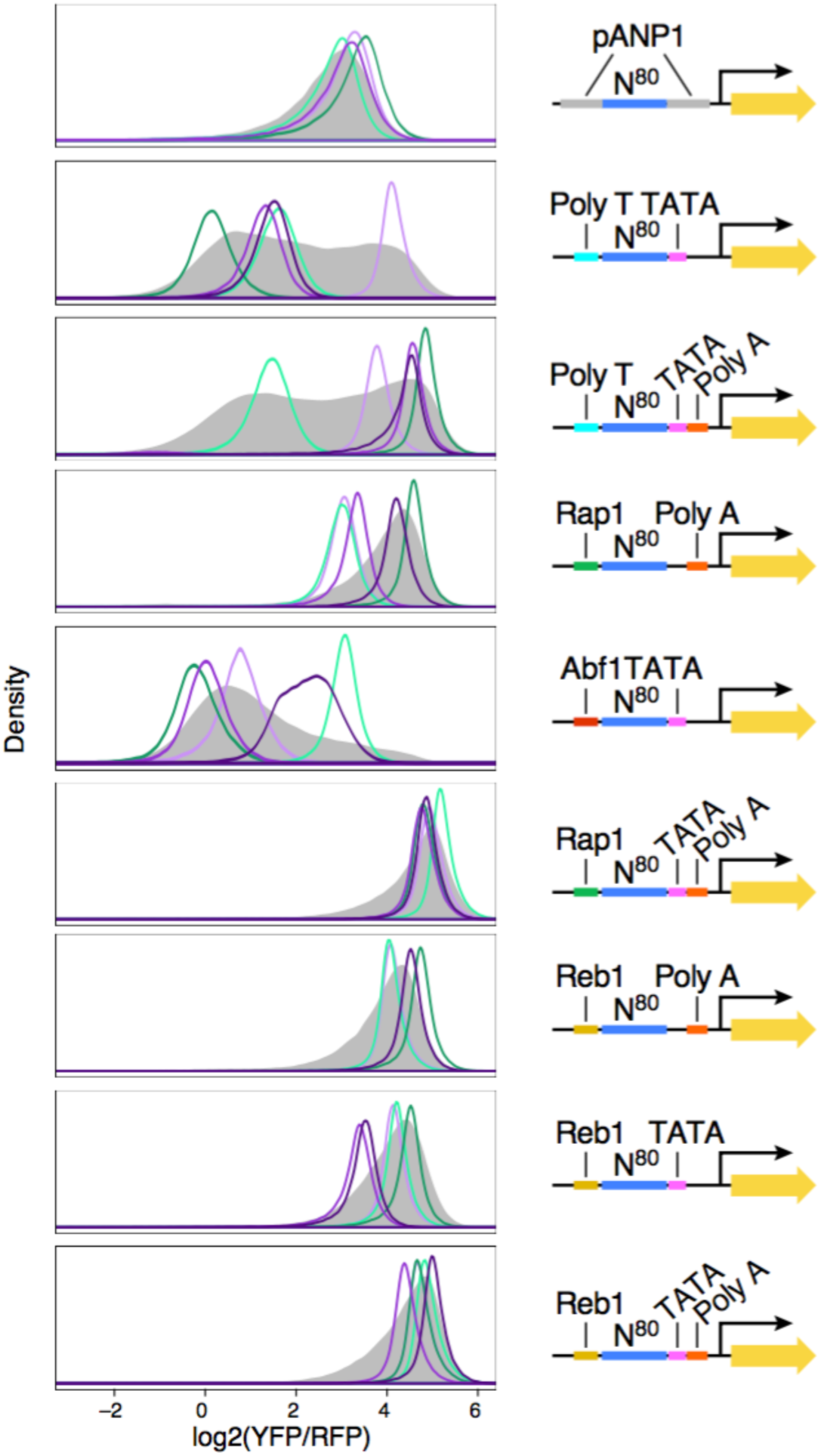
Random DNA yields diverse expression levels in all promoter scaffolds tested. For each promoter scaffold (right), shown are the distributions of expression levels (log_2_(YFP/RFP), *x* axis) measured by flow cytometry for the entire library (gray filled curves) and for a few selected clones, each from a different single promoter from the library (colored line curves).

**Supplemental Figure 2:**
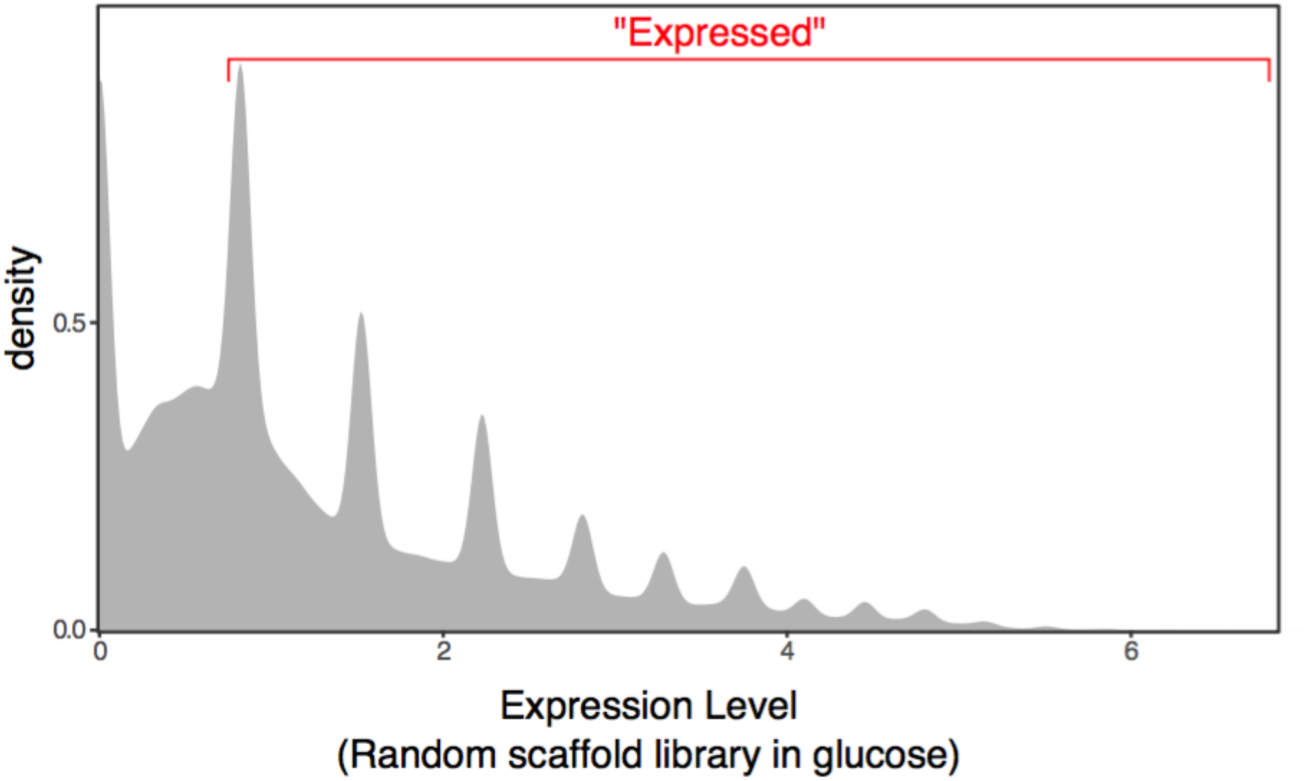
Expression level distribution of random DNA. Shown is the expression distribution of >2,000,000 promoters comprised of 3,811 random scaffolds, each in combination with ~660 random 80 bp oligos included in the middle. The periodic peaks occur at the expression bins and result from the large number of promoters that were only observed in a single bin (and so have a discrete expression level). We considered any promoters that, upon resorting, would end up in any of the non-zero bins as “expressed” (non-zero bins indicated in red; **Methods**). Note that the expression units are not equivalent to those used elsewhere; the dynamic range of this library is similar to that of the pTpA library, but contains fewer high-expressing promoters.

**Supplemental Figure 3:**
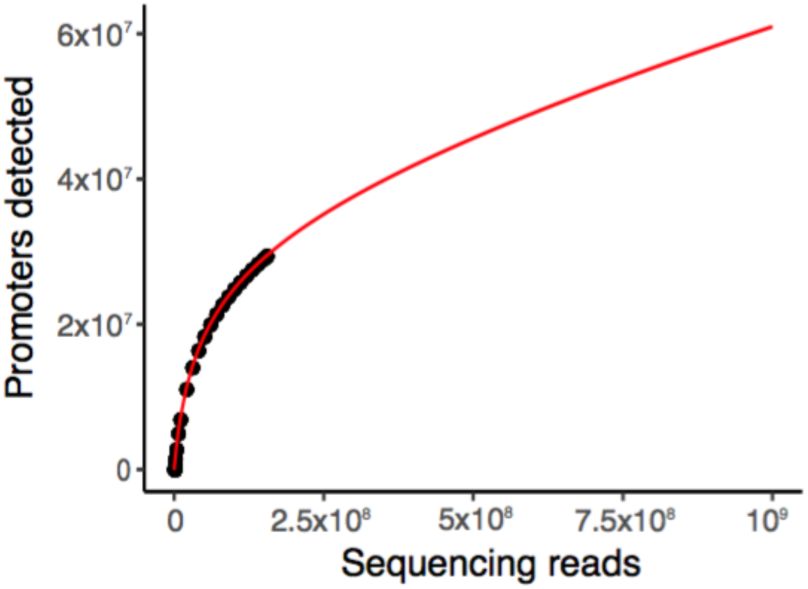
Saturation analysis of GPRA data. Shown are the numbers of distinct promoters detected when subsampling the pTpA+glucose sequencing data (black points), after combining reads from all expression bins. Red curve: promoters projected to be detected with additional sequencing (*54*).

**Supplemental Figure 4:**
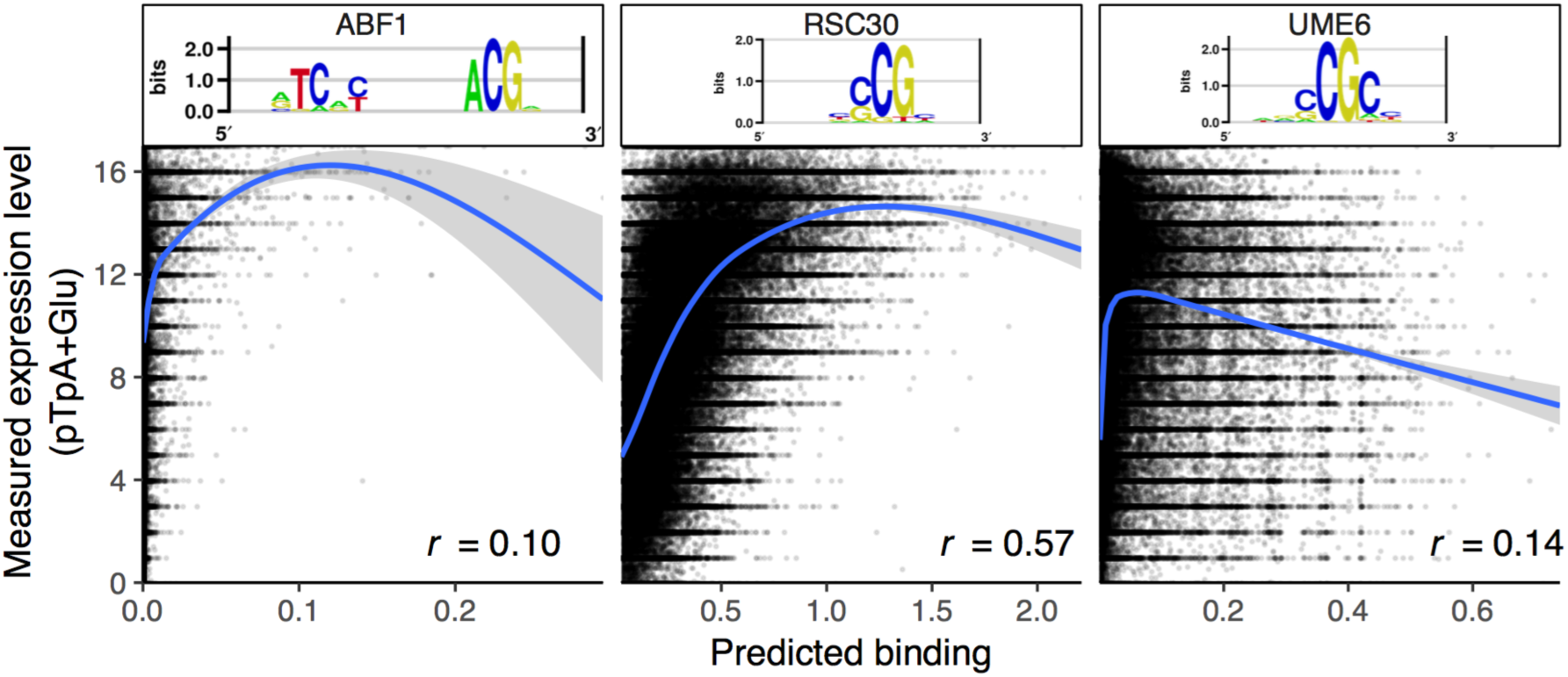
Relationship between predicted binding of individual TFs and expression level. Measured expression level (pTpA+Glu data; *y* axis) *vs*. predicted binding (*x* axis) for Abf1 (left), Rsc30 (middle), and Ume6 (right). Ume6 (a similar motif to Rsc30), is positively correlated with expression overall (*r*=0.14), but has a strong negative trend at high occupancies. Top: Motifs. Blue lines: GAM lines of best fit. Gray shaded areas: 95% confidence intervals.

**Supplemental Figure 5.**
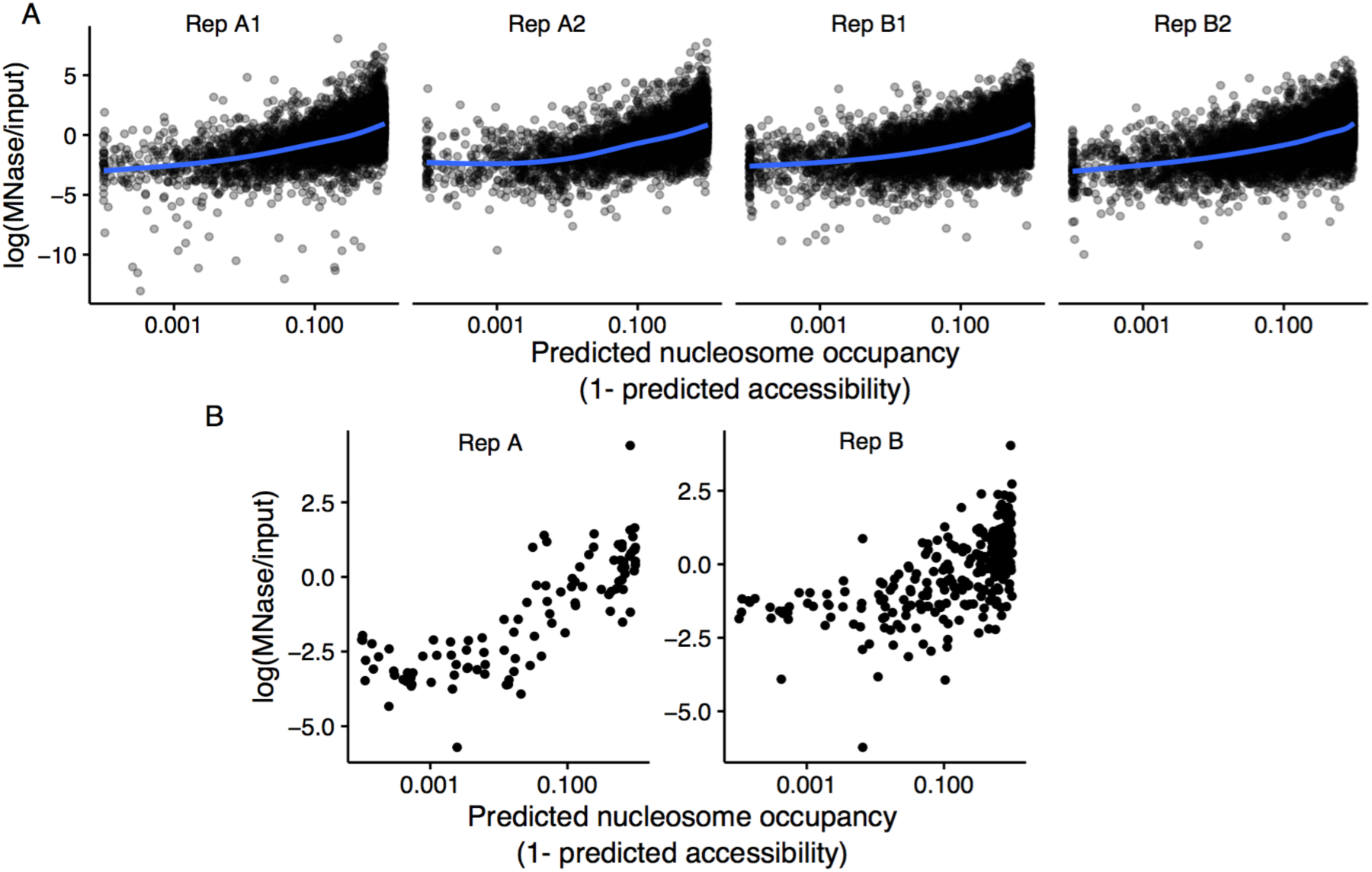
Predicted nucleosome occupancy. (**A**) Model predicted (*x* axis) *vs*. measured (MNase-Seq, *y* axis) nucleosome occupancy. Four MNase biological replicates are shown (**Methods**). (**B**) As in A, with replicates averaged, and only promoters present in both replicates shown.

**Supplemental Figure 6.**
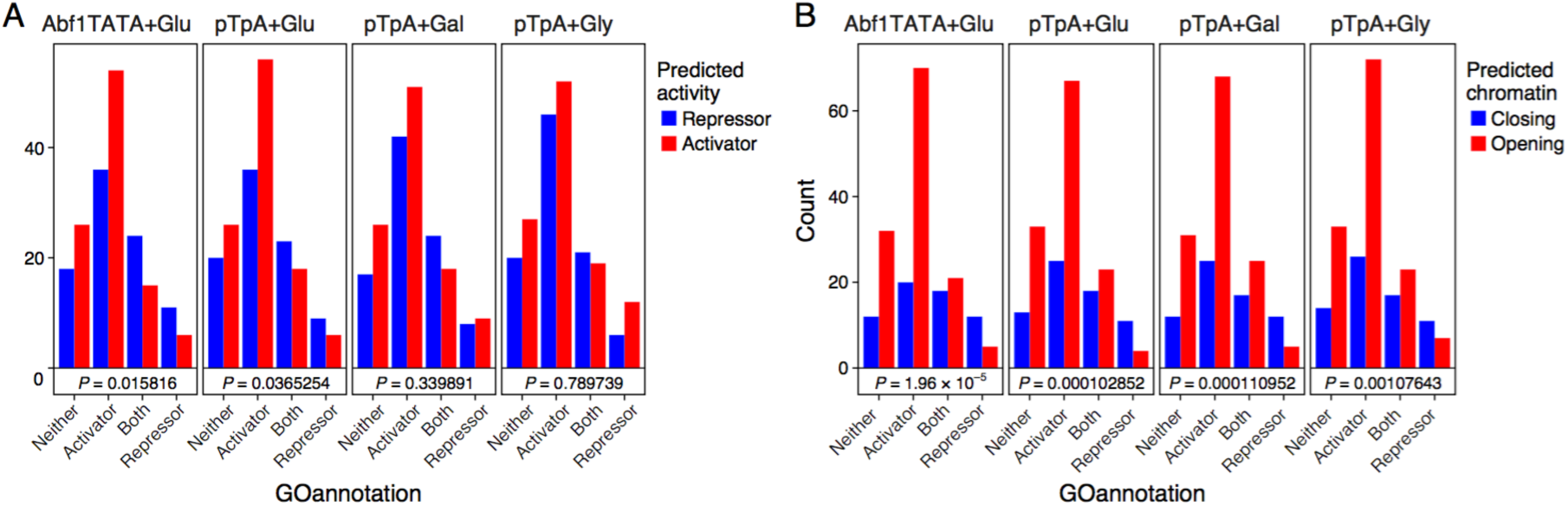
The billboard models identify biochemical activities of TFs. Shown are the number of TFs classified as activators, repressors, neither, or both in the yeast Gene Ontology (GO, **Methods**) (bars) and whether they are predicted as (**A**) repressor (blue) or activator (red); or (**B**) closing (blue) or opening (red) chromatin, by each model (label on top). Hypergeometric P-values for overlaps between predicted activator/repressor (or chromatin opener/closer), compared with activator/repressor GO annotations are as shown (“neither” and “both” categories are ignored).

**Supplemental Figure 7.**
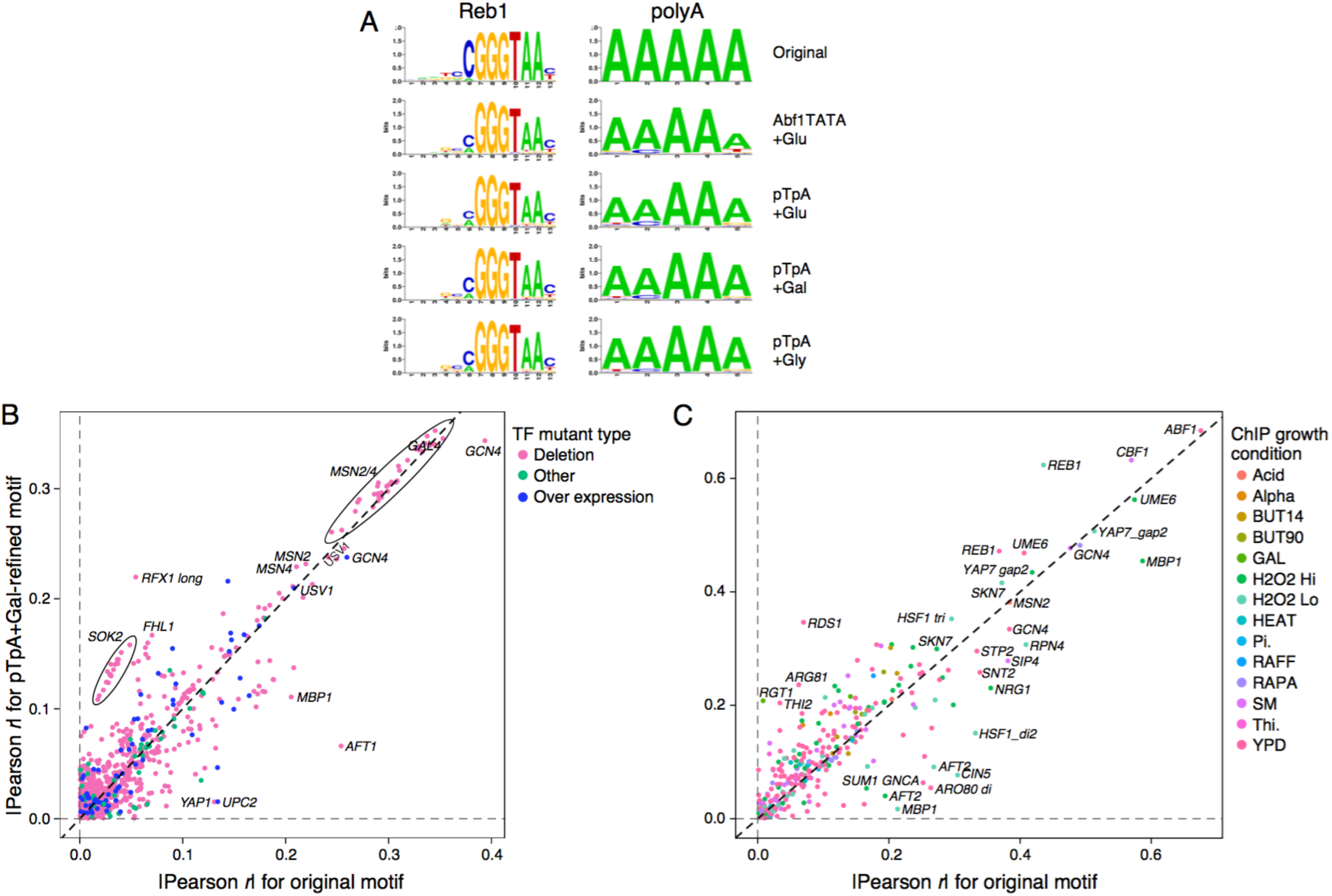
Model-refined motifs perform better in predicting TF binding and knockout effects in independent experiments. (**A**) Similar refinement in independent models. Comparison of the original TFBS motif (top) and model-refined motifs from each of the four models for two example motifs. (**B,C**) Shown are the absolute values of the Pearson correlation coefficient (|*r*|) when using either the original motifs (*x* axis) or the pTpA+Gal model-refined motifs (*y* axis) to predict whether (**B**) the gene’s expression will change in the corresponding TF mutant (compared to wild type (*21, 41*)) based on predicted binding to the promoter, or (**C**) a ChIP probe will be bound by the TF in a ChIP assay (*40*) based on predicted binding to the ChIP probe. (Here, data were not subsampled). Overall, model-refined motifs perform better (points above diagonal), but some perform worse. Reduced performance can be due to condition specific regulators that are minimally active in our tested growth conditions (*e.g.*, Gcn4), redundancy between motifs (*e.g.*, Hsf1 has mono-, di-, and trimeric motifs), and overfitting of the original motif to the test data (*e.g.*, ChIP-derived motifs tested on ChIP data).

**Supplemental Figure 8.**
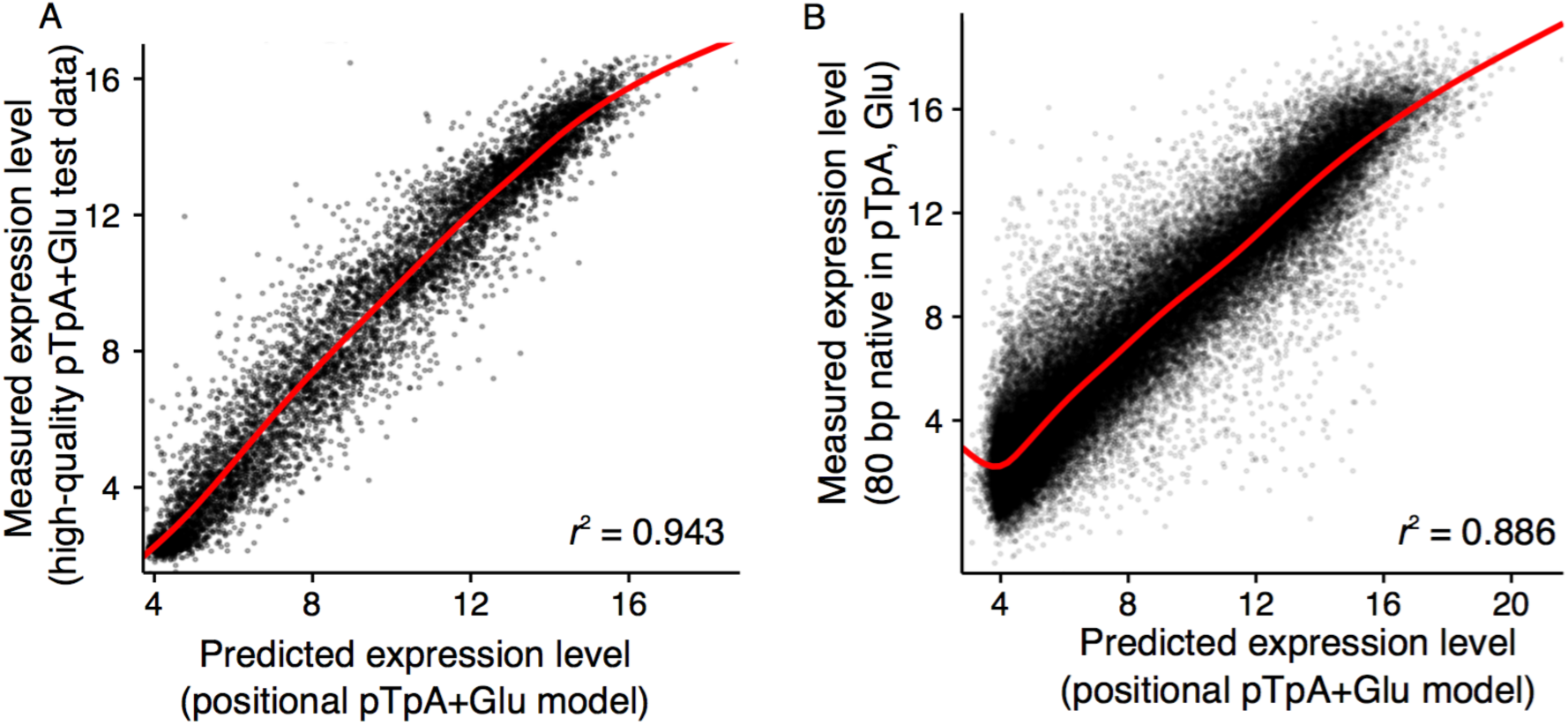
Positional models improve predictions. Position-specific pTpA+Glu model-predicted expression levels (*x* axes) *vs.* measured expression levels (*y* axes) for (**A**) high-quality test data in the pTpA promoter scaffold grown in glucose, and (**B**) native yeast promoter sequences, divided into 80 bp fragments and tested in the pTpA promoter scaffold grown in glucose.

**Supplemental Figure 9.**
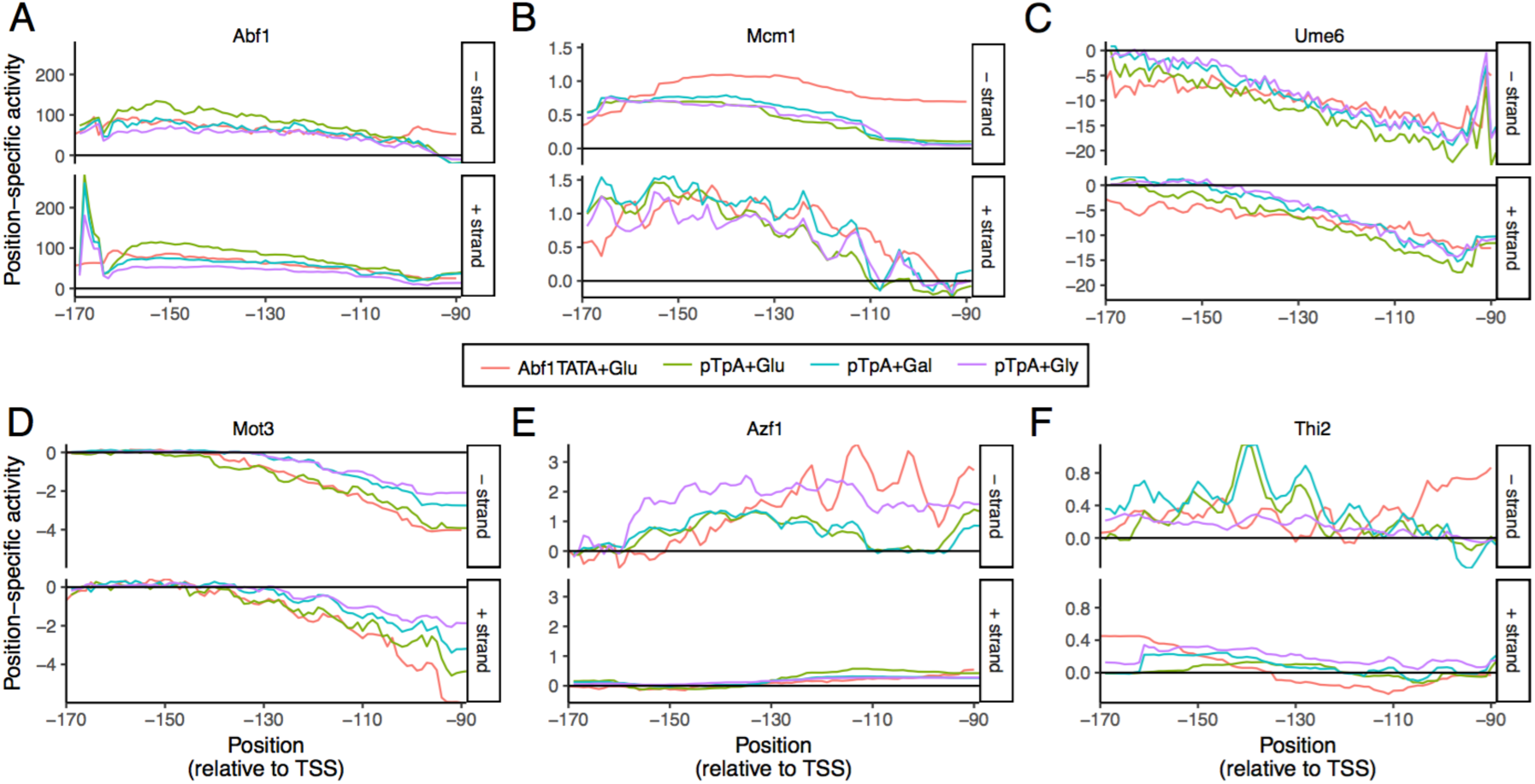
Positional preferences of TFs are prevalent and context-dependent. Position and strand preferences. Learned activity parameter values (*y* axis) for motifs in each position (*x* axis) and strand orientation (upper and lower panels) for each model (colors), for (**A**) Abf1, (**B**) Mcm1, (**C**) Ume6, (**D**) Mot3, (**E**) Azf1, and (**F**) Thi2.

**Supplemental Figure 10.**
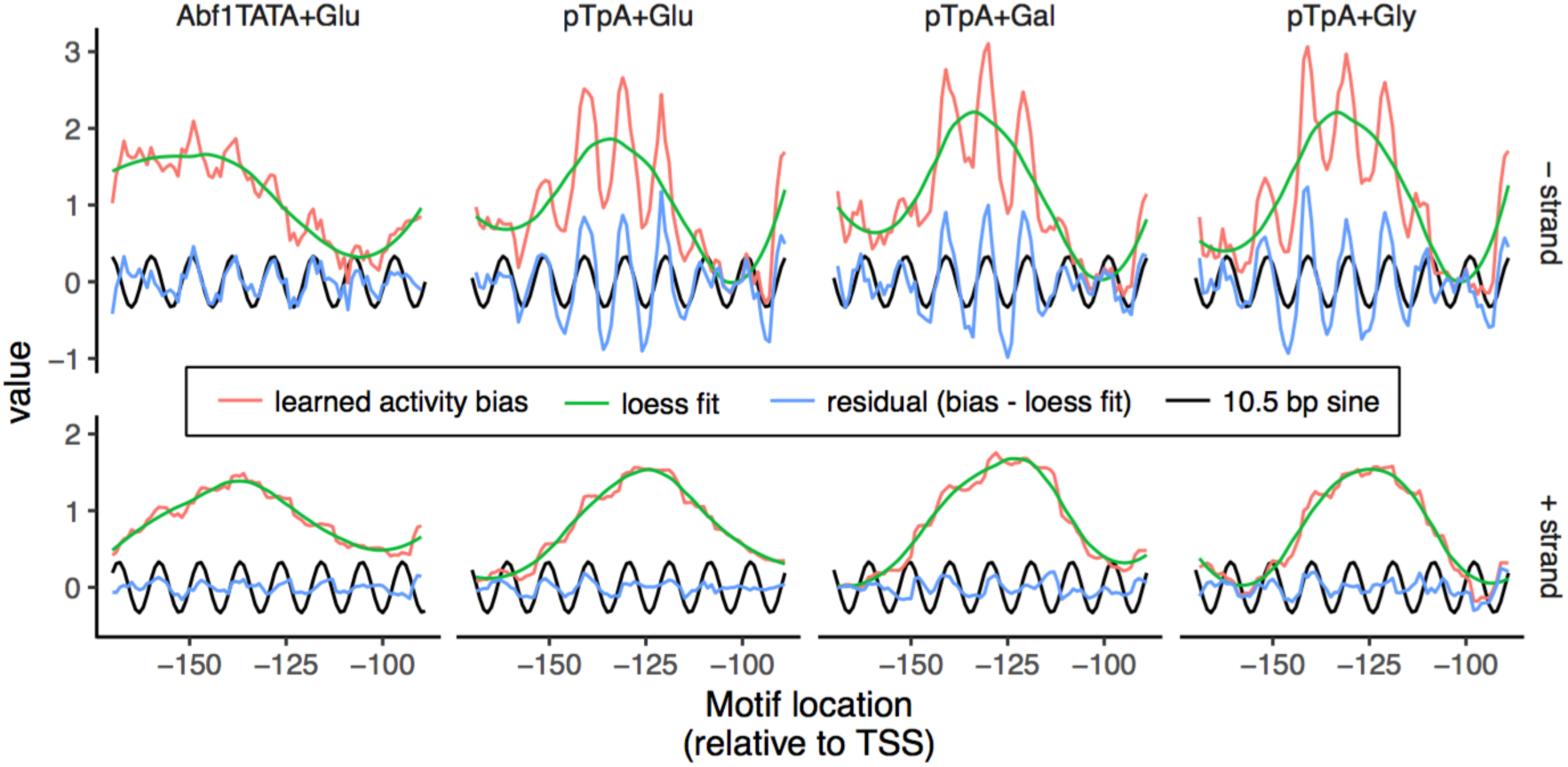
Capturing helically biased positional preferences. Plot shows, for each location within the promoter (*x* axis), the learned activity bias parameters (red curve; as in **Fig. 4b**) for the poly-A motif, long-range trend captured by a loess fit (green), and short-range residual activity bias after subtracting loess fit (blue) with reference 10.5 bp sine waves (black) for the minus strand (top) and plus strands (bottom) for the four different models (columns).

**Supplemental Figure 11.**
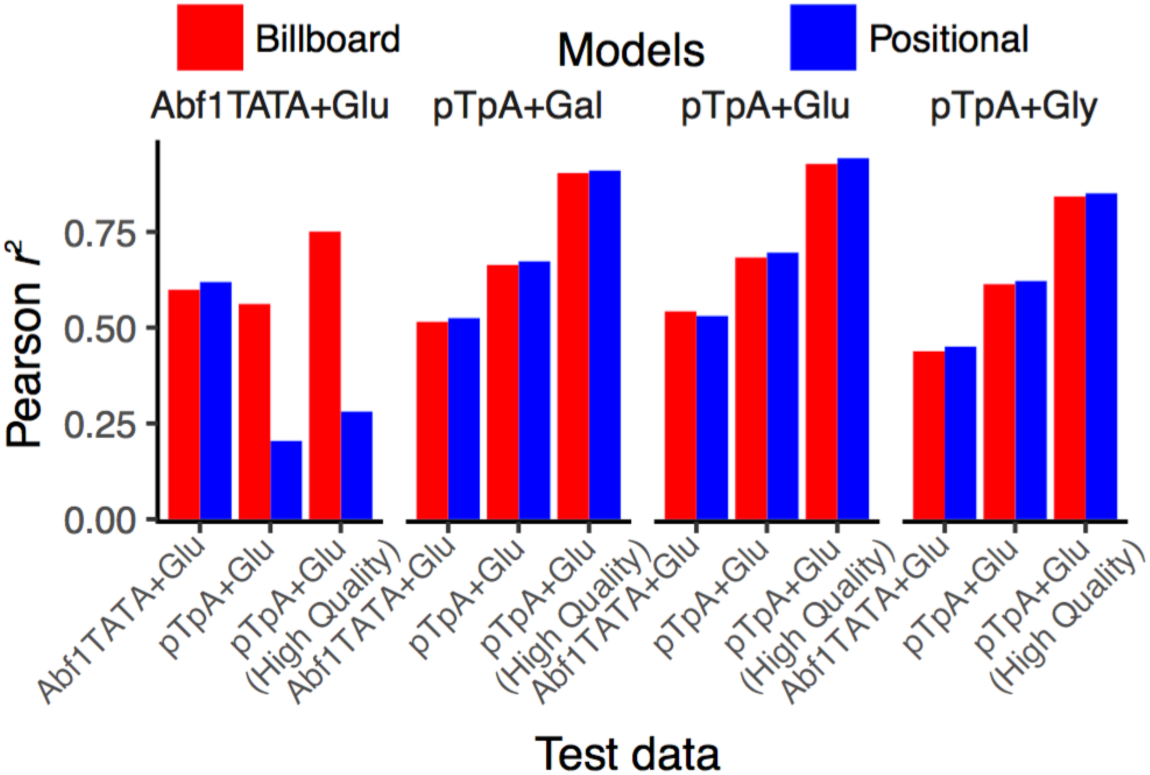
Positional preferences can be scaffold-specific. Modeling positional preferences increases predictive accuracy within the same scaffold but can drastically decrease it between scaffolds. For each training data set (four sub-panels) for both model types (colors), the Pearson *r*^2^ (*y* axis) capturing performance on each test dataset (*x* axis).

**Supplementary Table 1: Promoter scaffolds included in the scaffold library.** Sequences include 80 Ns in place of the random 80-mers and begin 13 bp upstream of the theoretical TSS.

**Supplementary Table 2: Motifs used in this study.** Motif IDs are from the YeTFaSCo database (*21*). Motifs excluded from the motif frequency analysis (**Fig. 1a**) are indicated.

